# Cytoplasmic innate immune sensing by the caspase-4 non-canonical inflammasome promotes cellular senescence

**DOI:** 10.1101/2020.10.16.342949

**Authors:** Irene Fernández-Duran, Núria Tarrats, Jodie Birch, Priya Hari, Fraser R. Millar, Morwenna Muir, Andrea Quintanilla, Valerie G. Brunton, João F. Passos, Juan Carlos Acosta

## Abstract

Cytoplasmic recognition of microbially derived lipopolysaccharides (LPS) in human cells is elicited by the inflammatory cysteine aspartic proteases caspase-4 and caspase-5, which activate non-canonical inflammasomes inducing a form of inflammatory programmed cell death termed pyroptosis. Here we show that LPS mediated activation of the non-canonical inflammasome also induces cellular senescence and the activation of tumour suppressor stress responses in human diploid fibroblasts. Interestingly, this LPS-induced senescence is dependent on caspase-4, the pyroptotic effector protein gasdermin-D and the tumour suppressor protein p53. Also, experiments with a catalytically deficient mutant suggest that caspase-4 proteolytic activity is not necessary for its role in senescence. Furthermore, we found that the caspase-4 non-canonical inflammasome is induced and assembled during Ras-mediated oncogene-induced senescence (OIS). Moreover, targeting caspase-4 in OIS showed that the non-canonical inflammasome is critical for SASP activation and contributes to reinforcing the cell cycle arrest in OIS. Finally, we observed that caspase-4 induction occurs *in vivo* in models of tumour suppression and ageing. Altogether, we are unveiling that cellular senescence is induced by cytoplasmic microbial LPS recognition by the caspase-4 non-canonical inflammasome and that this pathway is conserved in the senescence program induced by oncogenic stress.

## Introduction

Cellular senescence is a cell state characterized by a proliferative cellular arrest, a secretory phenotype, macromolecular damage and altered metabolism that can be triggered by several different stress mechanisms (Gorgoulis et al., 2019). Senescent cells produce and secrete a myriad of soluble and insoluble factors, including cytokines, chemokines, proteases and growth factors, collectively known as the senescence-associated secretory phenotype (SASP) (Acosta et al., 2008; Coppe et al., 2008; Kuilman et al., 2008) and interleukin-1 (IL-1) signalling is one of its critical signalling pathways (Acosta et al., 2013; Orjalo et al., 2009). The role of the SASP in cancer is complex and mechanistically ill-defined in ageing-associated diseases, two of the critical pathophysiological contexts where senescence is functionally relevant (Faget et al., 2019; McHugh and Gil, 2018). More recent evidence proposes that different triggers might induce distinctive SASP subsets with concrete functions (Herranz and Gil, 2018). Nonetheless, the SASP has started to incite interest as a potential therapeutic target in disease (Paez-Ribes et al., 2019; Soto-Gamez and Demaria, 2017). Therefore, a better understanding of the molecular machinery regulating the SASP is needed.

Pattern recognition receptors (PRRs) of the innate immune system are molecular sensors that are activated by microbial-derived pathogen-associated molecular patterns (PAMPs) or by damage-associated molecular patterns (DAMPs or alarmins) generated endogenously in cells under certain conditions of stress and damage (Takeuchi and Akira, 2010). Emerging data indicate a close relationship between these PRRs and cellular senescence. For example, the SASP is initiated following nucleic acid sensing by the cGAS-STING pathway and serum amyloid signalling through the PRR toll-like receptor-2 (TLR2) during oncogene-induce senescence is critical for the SASP and the cell cycle arrest (Dou et al., 2017; Gluck et al., 2017; Hari et al., 2019). Moreover, we have previously shown that inflammasomes are crucial for the SASP (Acosta et al., 2013). Inflammasomes are multiprotein platforms that induce the proteolytic activity of the inflammatory cysteine-aspartic protease caspase-1 which activates by proteolytic cleavage the proinflammatory cytokines IL-1β and interleukin-18 (IL-18). The canonical inflammasomes are assembled by PRRs of the nod-like receptor family, pyrin or by the cytoplasmic DNA sensor AIM2 (Lamkanfi and Dixit, 2014; Man and Kanneganti, 2016). Alternatively, the related inflammatory caspase-4 and caspase-5 (caspase-11 in mice) function as independent PRRs for cytoplasmic microbial lipopolysaccharide (LPS) activating a non-canonical inflammasome. Critically, activated non-canonical inflammasomes cleave the effector protein gasdermin-D, which induce a form of inflammatory programmed cell death termed pyroptosis (Kayagaki et al., 2015; Shi et al., 2015; Shi et al., 2014). Because the mechanism of SASP regulation by inflammasomes remain ill-defined, we decided to define the role of these inflammatory caspases in senescence. We show here that caspase-4 activation by cytoplasmic LPS triggers a senescence phenotype. Nonetheless, caspase-4 induction and non-canonical inflammasome assembly were observed in RAS^G12V^-oncogene-induced senescence (OIS). Moreover, we show here that the caspase-4 non-canonical inflammasome contributes critically to the establishment of the SASP and the reinforcement of the cell cycle arrest program during OIS, in a mechanism that is independent on its catalytic activity over its downstream pyroptotic target gasdermin-D. In all, we describe a new and critical function for cytoplasmic sensing by the caspase-4 non-canonical inflammasome in cellular senescence.

## Results

### Cytoplasmic LPS recognition by caspase-4 induces a senescent response in human diploid fibroblasts

While the canonical inflammasome can be activated with the microbial-derived molecule muramyl-dipeptide (MDP), the caspase-4 non-canonical inflammasome detects cytoplasmic bacterial LPS inducing gasdermin-D cleavage and pyroptosis. Gasdermin-D is the best characterized functional substrate of non-canonical inflammasomes, eliciting IL-1β secretion and pyroptotic cell death (He et al., 2015; Kayagaki et al., 2015; Shi et al., 2015). To compare the effect of activating the canonical and non-canonical inflammasome in senescence, we activated the caspase-1 or the caspase-4 inflammasomes by transfection of MDP or LPS in human IMR90 fibroblasts respectively. Similarly to other human cells expressing caspase-4 (Shi et al., 2014), IMR90 cells were sensitive to intracellular LPS in a dose-dependent manner (Figure S1A-C). In contrast, MDP transfection did not induce cell death (Figure S1A). LPS addition without further electroporation did not result in cell death, confirming the requirement of an intracellular location for LPS to trigger pyroptosis (Figure S1C). Knockdown of the pyroptosis effectors caspase-4 and gasdermin-D, but not caspase-1 impaired cell death mediated by LPS transfection, confirming that the cells were dying by pyroptosis (Figure S1D-F). Noticeably, cell death was detectable within the first hours after LPS transfection (Figure S1G). However, the fraction of cells surviving cell death after these hours remained stable and viable (Figure S1H), and this subpopulation was further examined.

Interestingly, this subset of cells displayed features of senescent cells such as decreased cell proliferation, increased levels of the cyclin-dependent kinase inhibitors p16^INK4a^ and p21^CIP1^ and increased senescence-associated-β-galactosidase (SA-β-galactosidase) activity (Figure S1I-J). In contrast, MDP transfection did not induce a senescent response (Figure S1J), indicating a specific role for the non-canonical inflammasome in PAMP-induced cellular senescence. Furthermore, the acquisition of a senescent phenotype was also accompanied by increased levels of caspase-4 (Figure S1J).

To examine the contribution of inflammatory caspases to the acquisition of a senescent phenotype following LPS transfection, we downregulated the expression of *CASP1* or *CASP4* with shRNA before transfection (Figure 1A). In contrast to caspase-1, caspase-4 was required for the acquisition of senescent features such as increased SA-β-galactosidase activity, decreased cell proliferation and induction of p21^CIP1^ and p16^INK4a^ in the subpopulation of cells surviving cell death (Figure 1B-D). Next, human *CASP1* and *CASP4* were ectopically expressed in IMR90 fibroblasts (Figure S2A). Overexpression of *CASP1* or *CASP4* alone resulted in a mild senescence induction with reduced cell proliferation and increased SA-β-galactosidase activity (Figure S2B-C). However, *CASP4* overexpression exacerbated the acquisition of senescent features in the presence of intracellular LPS (Figure 1E-F, S2D), indicating that caspase-4 expression is critical for non-canonical inflammasome induced senescence. Overall, these results suggest that the acquisition of a senescent phenotype following intracellular LPS exposure is mediated through caspase-4 in a dose-dependent manner.

**Figure 1.**
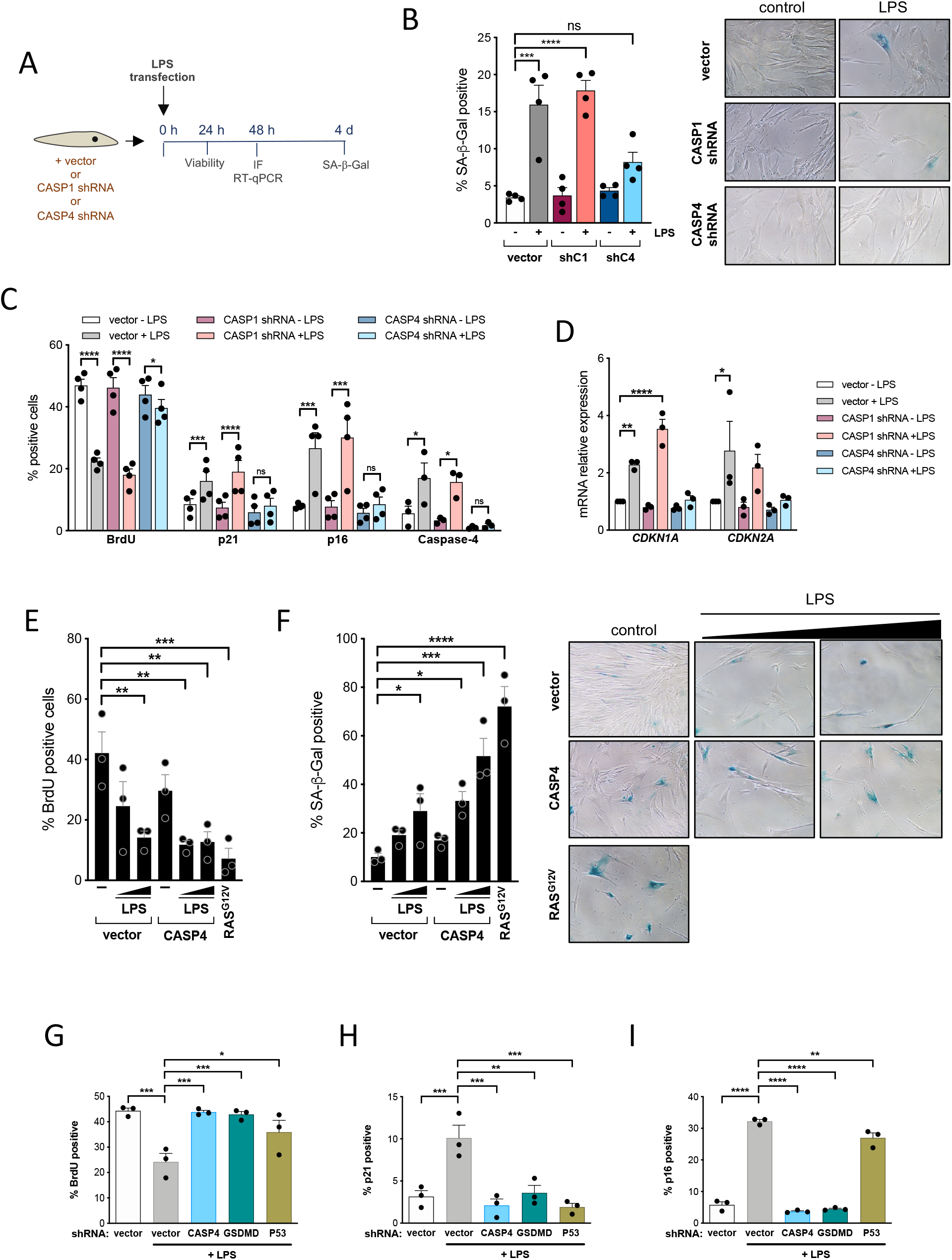
LPS-mediated caspase-4 activation induces a senescent phenotype in human primary fibroblasts. (A) Scheme of the experiment shown in B-D and Supplementary Figure D-E. IMR90 cells were infected with an empty pRS vector (vector) or a pRS vector targeting either *CASP1* (shC1) or *CASP4* (shC4) prior to transfection with 0.1 μg LPS / 5 × 10^^5^ cells. The acquisition of senescent features after LPS transfection was assessed by IF, RT-qPCR and SA-β-Gal activity. (B) SA-β-Gal activity was determined 4 days after transfection. Representative images for SA-β-Gal activity are shown. (C) BrdU incorporation and p16^INK4a^, p21^CIP1^ and caspase-4 levels of surviving cells were measured by IF 48 h after LPS transfection. (D) *CDKN1A* (p21^CIP1^) and *CDKN2A* (p16^INK4a^) mRNA relative expression was quantified by RT-qPCR 48 h after transfection. (E) *CASP4* was stably overexpressed prior to LPS transfection. Cells were transfected with increasing concentrations of LPS (0.1 or 1 μg LPS / 5 × 10^^5^ cells) and BrdU incorporation was measured 48 h after transfection. (F) Cells were treated as in (E) and SA-β-Gal activity was determined 4 days after transfection. Representative images for SA-β-Gal activity are shown. (G to I) IMR90 cells were stably infected with an empty pRS vector (vector) or a pRS vector targeting either *CASP4* (shC4), *GSDMD* (shGSDMD) or *TP53* (shP53) prior to transfection with 0.1 μg LPS / 5 × 10^^5^ cells. BrdU incorporation (G) and the levels of p21^CIP1^ (H) and p16^INK4a^ (I) were measured by IF 48 h after LPS transfection. Statistical significance in B, D-I was calculated using one-way analysis of variance (ANOVA). Statistical significance in C was calculated using two-tailed Student’s *t*-test. ****P < 0.0001, \*\*\**P* < 0.001, \*\**P* < 0.01, and \**P* < 0.05. ns, not significant. All error bars represent mean ± s.e.m; A represents 4 and B, D-I represents 3 independent experiments.

We decided to investigate the role of the caspase-4 substrate gasdermin-D and the critical senescence regulator p53 in LPS-induced senescence and pyroptosis by targeting their expression with shRNAs. In contrast to *GSDMD*, *TP53* knockdown failed to impair LPS mediated cell death, indicating a negligent role for p53 regulating caspase-4 dependent pyroptosis (Figure S1F). However, *TP53* and *GSDMD* knockdown significantly reduced LPS dependent cell growth arrest, and p53 and p21^CIP1^ induction (Figure 1G-H, S2E). Interestingly, *GSDMD* knockdown also had a strong effect on p16^INK4a^ and caspase-4 induction during LPS-induced senescence (Figure 1I, S2F). Altogether, these results indicate that cytoplasmic LPS sensing by the non-canonical inflammasome induced a senescence response that is dependent on caspase-4, gasdermin-D and p53 expression.

### The caspase-4 mediated LPS-induced senescent response is independent of IL-1β priming

Because overproduction of activated IL-1β can have detrimental effects (Afonina et al., 2015), the inflammasome activation is tightly regulated by a two-step mechanism. In some cases, a first signal, also called priming, is required to boost transcriptional levels of *IL1B*. The priming signal is then followed by a second signal which induces the assembly of the inflammasome (Lamkanfi and Dixit, 2014). Intriguingly, senescence induction by the sole overexpression of *CASP4* or *CASP1* in IMR90 fibroblasts did not induce transcriptional activation of *IL1B* (Figure 2A). In contrast, *IL1B* transcriptional levels were increased upon LPS transfection in a caspase-4 dependent fashion (Figure 2B). We have previously shown that TLR2 has a role in controlling *IL1B* expression and the SASP in cellular senescence (Hari et al., 2019). Thus, we decided to investigate if TLR2 mediated inflammasome priming could synergize with LPS-mediated caspase-4 induced senescence. As expected, the addition of the synthetic lipopeptides Pam2CSK4 and Pam3CSK4 (TLR2/6 and TLR1/2 agonists, respectively) but not LPS (TLR4 agonist) nor MDP to IMR90 cells highly induced *IL1B* mRNA levels (Figure S2G). Then, we observed that priming the inflammasome with the TLR2 agonist Pam2CKS4 significantly synergizes with LPS transfection to produce a robust IL-1β induction, and this induction was further enhanced by *CASP4* ectopic overexpression (Figure 2C). However, we did not observe a dramatic increase in cell cycle arrest or SA-β-galactosidase activity in LPS-induced senescence by addition of Pam2CKS4 (Figure 2D-E). Similar results were observed when TLR2 overexpressing cells were primed with the endogenous senescence-associated TLR2 alarmin A-SAA (Hari et al., 2019) or Pam2CKS4 with LPS transfection, which synergized in producing robust *IL1B* and SASP induction but without affecting dramatically LPS-induced cell cycle arrest or SA-β-Galactosidase activity (Figure S2H-L). However, the observed increase in *IL1B* mRNA levels in LPS transfected cells in all conditions is minimal if compared to the logarithmic increase observed in OIS (Figure 2C). These results suggest that additional signals to caspase-4 stimulation such as a sustained priming signalling are necessary for a robust *IL1B* and SASP induction during LPS-mediated cellular senescence. Moreover, these data suggest that the LPS-induced caspase-4 senescent response is independent of *IL1B* and the SASP.

**Figure 2.**
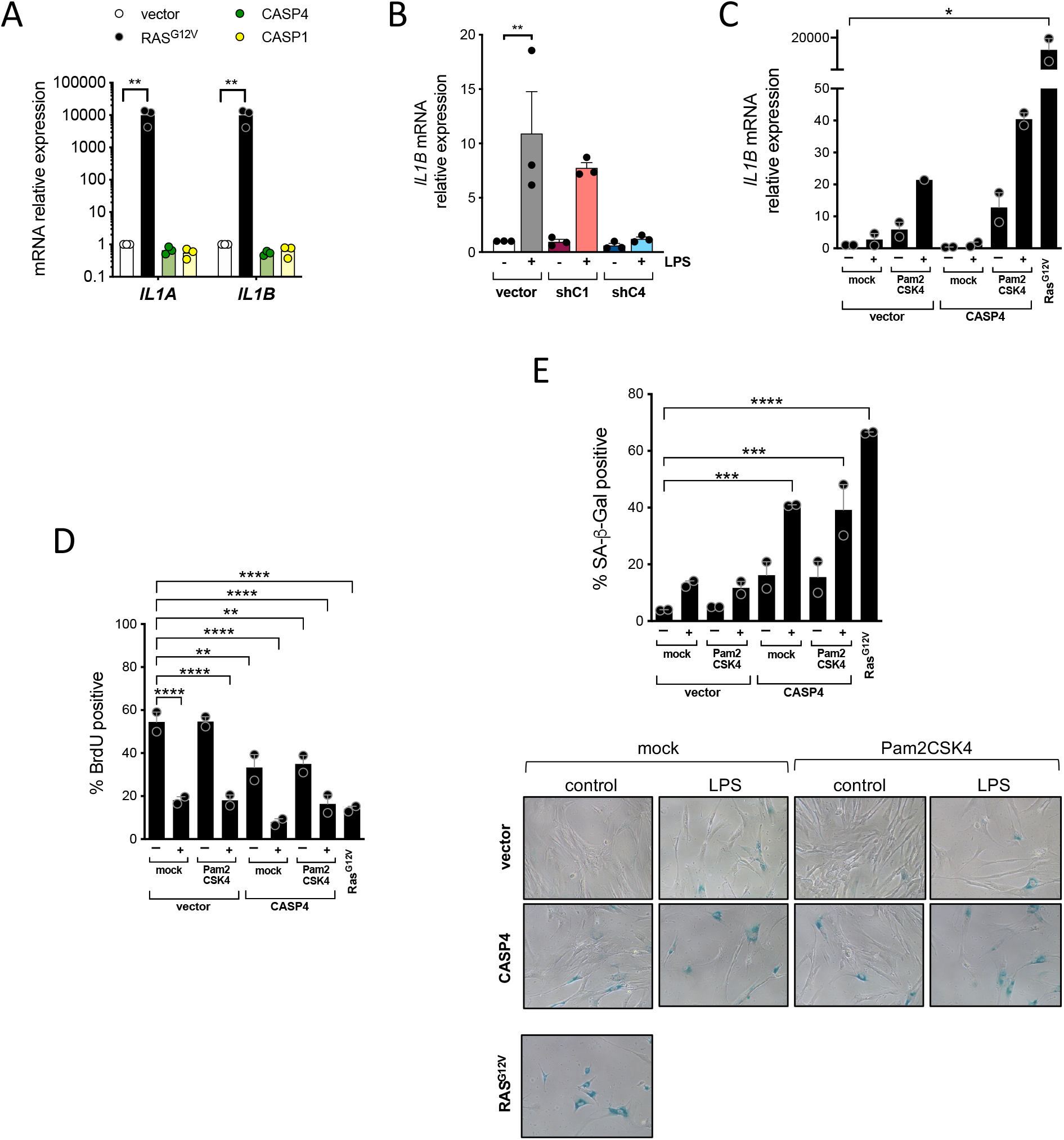
LPS-mediated caspase-4 induced senescence is independent on inflammasome priming. (A) *IL1B* mRNA relative expression levels were quantified by RT-qPCR in IMR90 infected with *CASP4*, *CASP1*, *RAS*^G12V^ expression vectors or empty vector (vector) control. (B) Cells were treated as shown in Figure 1A. *IL1B* mRNA relative expression was quantified by RT-qPCR 48 h after LPS transfection. (C-E) IMR90 cells were infected with *CASP4* or *RAS*^G12V^ expression vectors or empty vector (vector) control. After 3 h treatment with Pam2CSK4, cells were transfected with LPS (0.1 μg LPS / 5 × 10^^5^ cells). *IL1B* mRNA relative expression (C) and BrdU incorporation (D) were measured by IF and RT-qPCR respectively 48 h after LPS transfection. (E) SA-β-Gal activity was determined 4 days after LPS transfection. Representative images are shown. For A and B, sstatistical significance was calculated using one-way analysis of variance (ANOVA). ****P < 0.0001, \*\*\**P* < 0.001, \*\**P* < 0.01, and \**P* < 0.05. All error bars represent mean ± s.e.m of 3 independent experiments. For C-E, sstatistical significance was calculated using one-way analysis of variance (ANOVA). ****P < 0.0001, \*\*\**P* < 0.001, \*\**P* < 0.01, and \**P* < 0.05. Error bars represent the average ± range of 2 representative experiments.

### The caspase-4 proteolytic activity is not necessary for LPS-induced senescence

The active site of human caspase-4 has been well characterized and is associated to the residue C258 (Faucheu et al., 1995), and point mutations of this amino acid renders the protein catalytically inactive (Shi et al., 2014; Sollberger et al., 2012). To further study the contribution of the protease activity of caspase-4 protease to senescence, *CASP4* wild-type and the catalytically death mutant C258A (*CASP4*^C258A^) were overexpressed in IMR90 cells, and the phenotypical outcomes were assessed. Overexpression of wild-type *CASP4* or *CASP4*^*C258A*^ reduced cellular proliferation and increased SA-β-galactosidase activity to a similar extent (Figure 3A-B). Next, *CASP4* wild-type and *CASP4*^*C258A*^ were stably overexpressed in IMR90 fibroblasts before LPS transfection (Figure 3C). *CASP4*^*C258A*^ but not *CASP4* wild-type expressing IMR90 cells were resistant to cell death after LPS transfection (Figure 3D, S3A), indicating a dominant negative role for the caspase-4 defective form in pyroptosis. However, both *CASP4* wild-type and *CASP4*^*C258A*^ overexpressing cells remained equally sensitive to the acquisition of senescent features after LPS challenge (Figure 3E-F, S3B). These results suggest that, in contrast to caspase-4 mediated pyroptosis, the role of caspase-4 in LPS-induced senescence is independent of its catalytic activity.

**Figure 3.**
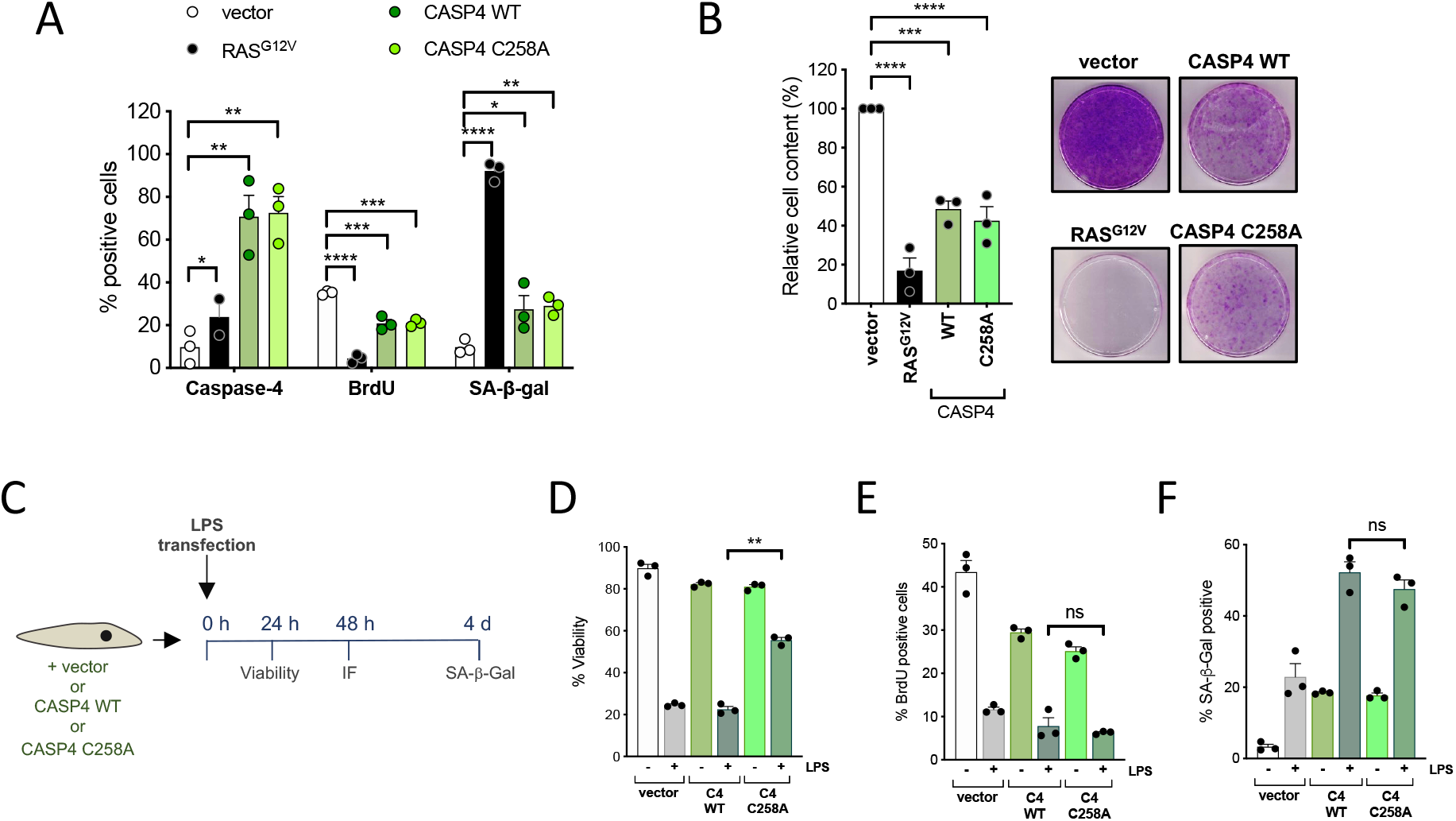
Caspase-4 mediated regulation of senescence is independent of its catalytical function. (A-B) IMR90 cells were infected with wild-type (WT) *CASP4*, catalytically inactive (C258A) *CASP4* or the empty vector (vector). Overexpression of *RAS*^G12V^ was used as a positive control for the induction of senescence. (A) Caspase-4 levels, BrdU incorporation and SA-β-galactosidase activity were measured 4 days after equal number of cells were seeded. (B) Relative cell content (left) was quantified 15 days after equal number of cells were seeded; representative images (right) of crystal violet stained cells are shown. (C) Scheme of the experiment shown in D-F and Supplementary Figure A-B. IMR90 cells were infected with wild-type (WT) *CASP4*, catalytically inactive (C258A) *CASP4* or the empty vector (vector) prior to transfection with 0.1 μg LPS / 5 × 10^^5^ cells. (D) Cell viability was measured 24 h after transfection. (E) BrdU incorporation was measured 48 h after transfection. (F) SA-β-Gal activity was determined 4 days after transfection. Statistical significance in A and B was calculated using one-way analysis of variance (ANOVA). Statistical significance in D-F was calculated using two-tailed Student’s *t*-test. ****P < 0.0001, \*\*\**P* < 0.001, \*\**P* < 0.01, and \**P* < 0.05. ns, not significant. All error bars represent mean ± s.e.m of 3 independent experiments.

### The caspase-4 non-canonical inflammasome is induced and assembled during oncogene-induced senescence

Next, the role of inflammatory caspases in oncogene-induced senescence (OIS) was investigated. To induce OIS, *HRAS*^*G12V*^ (hereafter *RAS*^*G12V*^) was constitutively overexpressed in IMR90 human fibroblasts. *RAS*^*G12V*^ overexpression reduced cellular proliferation and increased SA-β-galactosidase activity (Figure 4A). Coinciding with the upregulation of the cell cycle inhibitors p21^CIP1^, p16^INK4a^ and p15^INK4b^, caspase-4 expression was increased both at the mRNA and protein levels upon *RAS*^*G12V*^ overexpression (Figure S4A-B, 4B-C). Next, we took advantage of an inducible system extensively employed by us and others to exert tight control over the onset of senescence (Boumendil et al., 2019). In this system, a mutant form of the estrogen receptor (ER) ligand-binding domain is fused to the protein of interest (RAS); consequently, ER:RAS cells undergo OIS after addition of 4-hydroxytamoxifen (4OHT) (Figure 4D). As expected, IMR90 ER:RAS cells underwent cell proliferation arrest and showed increased SA-β-galactosidase activity compared to non-treated IMR90 ER:RAS upon 4OHT addition (Figure 4D). A time-course experiment using this system revealed that *CASP4* mRNA levels increase in parallel to the exponential increase in *IL1B* mRNA expression in cells undergoing OIS (Figure 4E-F), and in caspase-4 protein (Figure 4G). Also, we observed caspase-4 induction in paracrine senescence, and senescence induced by DNA damage with etoposide (Figure S4C-D). Oligomerization of caspase-4 protein is essential for its activity (Shi et al., 2014). These caspase-4 oligomers can be revealed in lysates crosslinked with disuccinimidyl suberate (DSS) as high weight migratory bands in a western blot (Choi et al., 2019). Endogenous caspase-4 oligomerization was detected in IMR90 ER:RAS senescent cells but not in control cells from 3 days after 4OHT treatment (Figure 4H, S4E). Moreover, caspase-4 proteolytic activity was also increased in IMR90 ER:RAS cells 4 and 8 days after the addition of 4OHT (Figure 4I). Altogether these results demonstrate that caspase-4 expression is induced and the non-canonical inflammasome is assembled in OIS.

**Figure 4.**
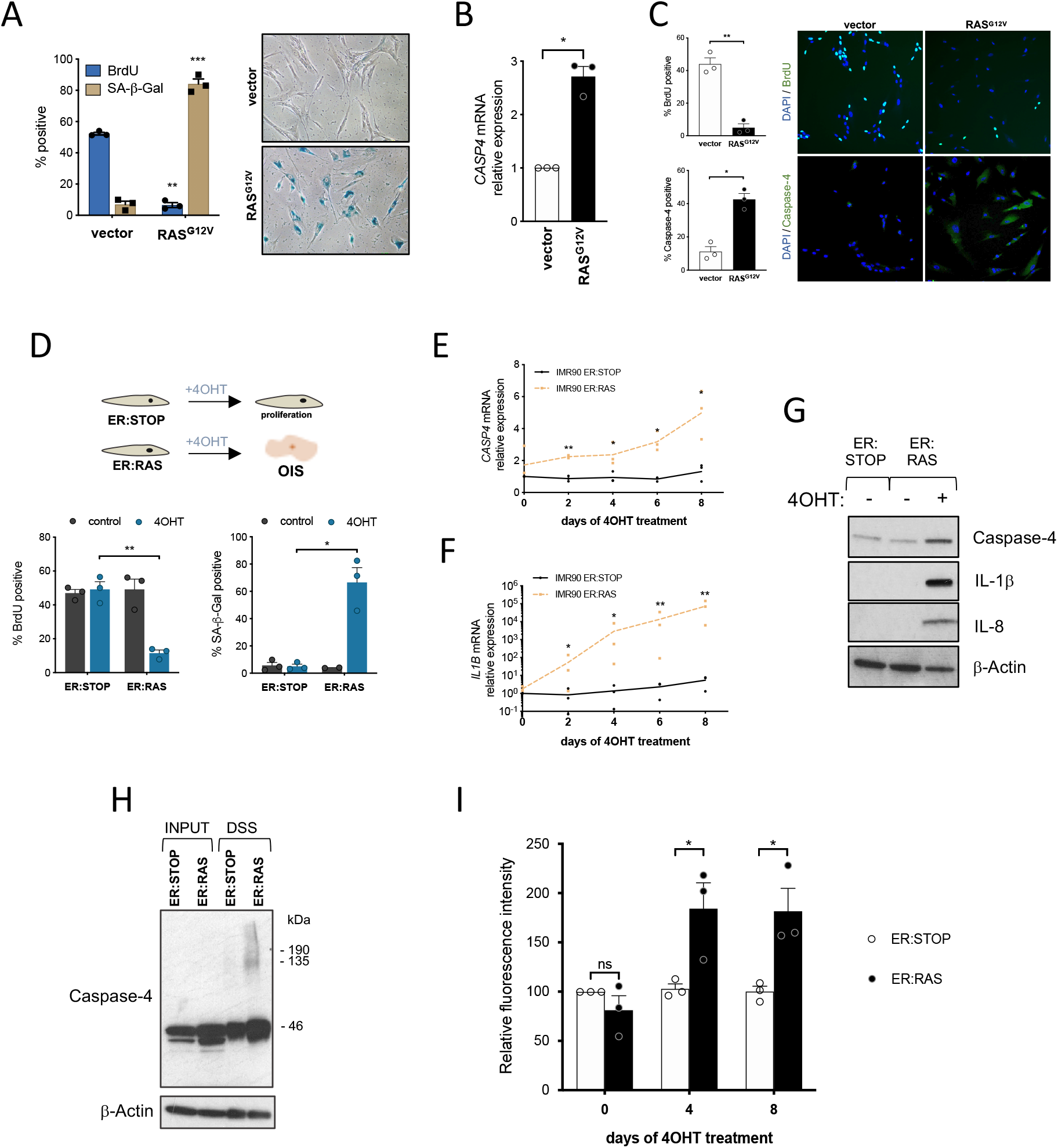
The caspase-4 non-canonical inflammasome is activated in Oncogene-induced senescence. (A) IMR90 cells were infected with *RAS*^G12V^ expression vector to induce OIS or empty vector (vector). BrdU incorporation and SA-β-Gal activity were measured 4 days after equal number of cells were seeded (left). Representative images (right) for SA-β-Gal activity are shown. (B) *CASP4* mRNA relative expression was quantified in IMR90 cells undergoing *RAS*^G12V^ after infection with *RAS*^G12V^ expression vector- (C) IMR90 cells were infected with *RAS*^G12V^ expression vector or empty vector (vector). BrdU incorporation and caspase-4 levels were measured by IF in RAS^G12V-^OIS and control cells 4 days after equal number of cells were seeded. (D) IMR90 cells were infected with a control (ER:STOP) or an ER:RAS vector. Upon addition of 4OHT, ER:RAS cells undergo OIS. (E) Time-course experiment of *CASP4* mRNA relative expression in IMR90 ER:STOP and ER:RAS cells treated with 4OHT for 0, 2, 4, 6 and 8 days. (F) Time-course experiment of *IL1B* mRNA relative expression in IMR90 ER:STOP and ER:RAS cells treated with 4OHT for 0, 2, 4, 6 and 8 days. (G) IMR90 ER:STOP and ER:RAS were treated or not with 4OHT during 8 days. Caspase-4, IL-1β and IL-8 levels were analyzed by immunoblotting. (H) IMR90 ER:STOP and ER:RAS cells were treated with 4OHT for five days, then cells were collected and subjected to disuccinimidyl suberate (DSS) crosslinking. After SDS-PAGE separation, both DSS-crosslinked samples and inputs were probed for caspase-4 by immunoblotting. (I) IMR90 ER:STOP and ER:RAS cells were treated with 4OHT and LEVD-AFC cleavage was measured in low serum (0.5% FBS) cultured cells 0, 4 and 8 days after 4OHT addition. All statistical significance was calculated using using two-tailed Student’s *t*-test. \*\*\**P* < 0.001, \*\**P* < 0.01, and \**P* < 0.05 and ns, not significant. All error bars represent mean ± s.e.m of 3 independent experiments.

### The caspase-4 non-canonical inflammasome is required for inflammatory signalling in OIS

Next, a functional role for caspase-4 in OIS was investigated in our IMR90 ER:RAS system. Global changes in mRNA expression upon *CASP4* depletion were analyzed. IMR90 ER:STOP and ER:RAS cells were transfected with control and *CASP4*-targeting small-interfering RNA (siRNA), samples were collected 5 and 8 days after 4OHT addition and subjected to transcriptomic analysis (Figure 5A). Knockdown control of the experiment was performed by mRNA expression analysis for *CASP4*, showing a substantial reduction of its expression upon siRNA targeting at both time points (Figure S5A). Furthermore, *CASP4* was the top downregulated gene in *CASP4* siRNA-targeted compared to non-target siRNA control RAS^G12V^ cells both 5 and 8 days after the addition of 4OHT (Figure S5B). Similarities between replicates and differences between conditions were confirmed by principal component analysis visualization and heatmap sample clustering (Figure S5C-D). Differentially expressed gene analysis identified 557 and 478 genes significantly differentially expressed (FDR 10%) upon *CASP4* knockdown, of which 340 and 240 were induced in a *CASP4* dependent fashion in *RAS*^*G12V*^-OIS cells 5 and 8 days after 4OHT addition respectively (Figure 5A). Gene set enrichment analysis (GSEA) of 50 hallmark gene sets of the transcriptomic data showed a CASP4 dependent regulation in *RAS*^*G12V*^-OIS cells of gene sets related to inflammatory processes, including TNF-α signalling and interferon responses, both 5 and 8 days after the addition of 4OHT (Figure 5B). Enrichment plots of the gene signature hallmark “INFLAMMATORY RESPONSE” showed a positive correlation of this gene set expression with *CASP4* (Figure S5E). Plotting a heatmap of the fold change values of control IMR90 ER:STOP and *CASP4* knockdown vs control IMR90 ER:RAS cells of all genes included in the “INFLAMMATORY RESPONSE” gene set revealed a pattern by which the increased expression of inflammatory-related genes in senescence is abrogated if *CASP4* is targeted, including SASP factors (Figure 5C). Changes in the expression of *IL1A* and *IL1B* were validated by RT-qPCR (Figure 5D-E). The serum amyloid A (SAA) proteins SAA1 and SAA2 belong to a family of apolipoproteins known to activate innate and adaptive immune cells and have recently been identified as SASP factors (Hari et al., 2019). Of note, the expression of *SAA1* and *SAA2* was also decreased when *CASP4* was targeted in OIS (Figure 5F-G). Targeting *CASP4* also reduced the amount of intracellular IL-1α, IL-1β, IL-6 and IL-8 protein to a similar extent than *CASP1* targeting (Figure S5F). Moreover, the levels of intracellular mature IL-1β were also significantly and similarly reduced when either *CASP1* or *CASP4* were targeted (Figure 5H). Furthermore, the concentration of secreted IL-1β was significantly reduced in conditioned media of RAS induced cell cultures when *CASP4* was targeted (Figure 5I). Overall, these results suggest that caspase-4 upstream of caspase-1 are required for a full SASP activation in OIS.

**Figure 5.**
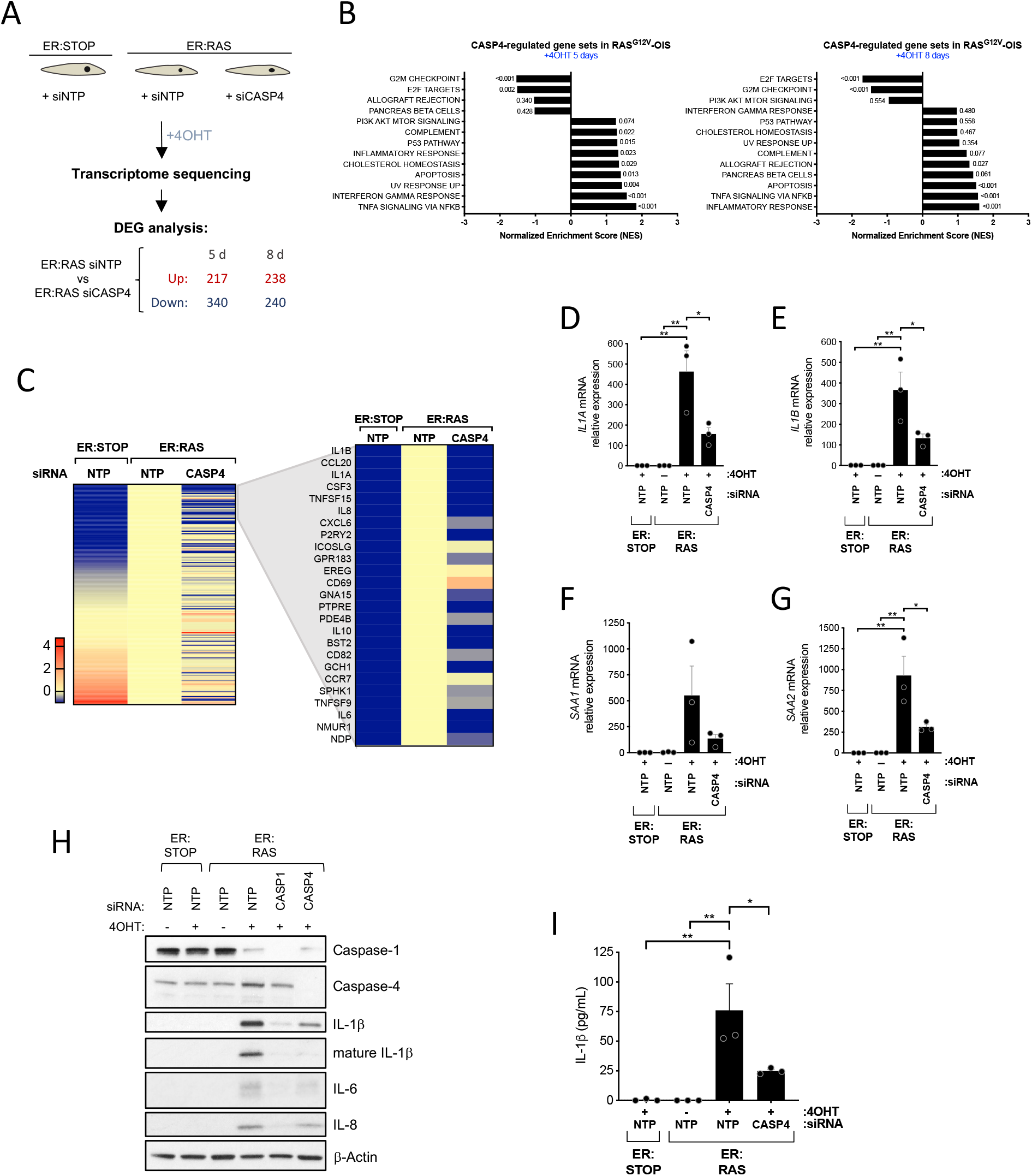
Caspase-4 activation controls the proinflammatory SASP. (A) Schematic diagram of the experimental approach. ER:STOP and ER:RAS IMR90 cells were targeted with either control (non-targeting pool, NTP) or *CASP4*-targeting siRNA. All cells were treated with 4OHT from day 0. RNA was extracted at day 5 and 8 after the addition of 4OHT and subjected to transcriptomic analysis. Differentially expressed gene (DEG) analysis was performed and the number of significant upregulated and downregulated genes 5 and 8 after the addition of 4OHT upon *CASP4*-targeting in ER:RAS cells is shown. (B) Normalized Enriched Scores (NES) of a set of 50 curated hallmark gene signatures were calculated based on the DEG analysis performed between control and *CASP4*-knockdown ER:RAS samples after 5 and 8 days of 4OHT treatment. Gene sets with a false discovery rate (FDR) q-value of ≤ 0.25 at least in one of the timepoints are shown. P-values for each gene set are indicated next to the corresponding bar. (C) Heatmap of the log2FC values of all 175 genes included in the “INFLAMMATORY RESPONSE” GSEA gene set of control ER:STOP and *CASP4*-knockdown ER:RAS compared to control ER:RAS after 5 days of 4OHT treatment. The top 25 differentially expressed signature genes in *RAS*^G12V^-OIS are zoomed in. (D) *IL1A* mRNA relative expression levels were quantified by RT-qPCR after 5 days of 4OHT treatment in ER:STOP and ER:RAS cells transfected with the indicated siRNA. (E) *IL1B* mRNA relative expression levels were quantified by RT-qPCR after 5 days of 4OHT treatment in ER:STOP and ER:RAS cells transfected with the indicated siRNA. (F) *SAA1* mRNA relative expression levels were quantified by RT-qPCR after 8 days of 4OHT treatment in ER:STOP and ER:RAS cells transfected with the indicated siRNA. (G) *SAA2* mRNA relative expression levels were quantified by RT-qPCR after 8 days of 4OHT treatment in ER:STOP and ER:RAS cells transfected with the indicated siRNA. (H) IMR90 ER:STOP/ER:RAS cells were transfected with control (NTP), *CASP1* or *CASP4-*targeting siRNA and treated with 4OHT or not during 8 days as indicated. Lysates were subjected to immunoblotting analyses with the indicated antibodies. (I) IMR90 ER:STOP/ER:RAS cells were treated or not 8 days with 4OHT as indicated and secreted IL-1β was quantified by ELISA. Statistical significance in A-E was calculated using one-way analysis of variance (ANOVA). \*\**P* < 0.01, and \**P* < 0.05. All error bars represent mean ± s.e.m of 3 independent experiments.

The SASP can induce senescence in adjacent growing cells through paracrine signaling, which is dependent on IL-1 signaling (Acosta et al., 2013). Because *CASP1* and *CASP4*-targeting reduced the production of several SASP factors and, in particular, limited the secretion of IL-1β, we next examined whether inflammatory caspases are implicated in SASP induction during paracrine senescence (Figure S5G). Conditioned media from IMR90-ER:RAS senescent cells added to growing IMR90 fibroblasts produced the induction of *IL1A, IL1B*, *IL8* and *IL6*, which was impaired when *CASP4* was targeted (Figure S5G). In contrast, *CASP1* targeting did not affect the induction of the paracrine SASP (Figure S5G). Overall, these data suggest that the caspase-4 non-canonical inflammasome controls SASP activation during paracrine senescence.

We then investigated the role of gasdermin-D in OIS. While mRNA levels remained unaffected (Figure S6A), gasdermin-D was found to be cleaved during OIS (Figure S6B). However, in contrast to *CASP1* or *CASP4*-targeting, *GSDMD* knockdown did not impair *IL1A*, *IL1B*, *IL8* or *IL6* mRNA induction (Figure S6C-D). Moreover, whereas targeting either *CASP1* or *CASP4* resulted in a significantly lower concentration of IL-1β in conditioned media from OIS cells, *GSDMD* knockdown did not alter IL-1β secretion (Figure S6E), suggesting that IL-1β secretion is dependent on caspase-1 and caspase-4 but independent on gasdermin-D in OIS. Altogether, these results indicate that the caspase-4 non-canonical inflammasome is a critical regulator of the SASP in OIS.

### The caspase-4 non-canonical inflammasome contributes to the cell cycle arrest in OIS

To investigate the role of the non-canonical inflammasome in regulating the cell cycle arrest program in OIS, we targeted *CASP4* using RNAi (Figure S6F). *CASP4*-targeting significantly rescued early proliferation arrest during OIS (Figure 6A) and increased the total cell content in a long-term cell growth assay (Figure 6B). Moreover, targeting *CASP4* during OIS modestly but significantly decreased SA-β-galactosidase activity (Figure S6G-H). A GSEA of 50 hallmark gene sets showed a negative regulation of *CASP4* in *RAS*^G12V^-OIS of the gene signatures “G2M CHECKPOINT” and “E2F TARGETS”, and the expression of the *CDKN2A* (p16^INK4a^-p14^ARF^) and *CDKN2B* (p15^INK4b^) locus, but not p21^CIP1^ (Figure 5B, 6C, S6H). Indeed, targeting *CASP4* reduced p16^INK4a^ expression and rescued the phosphorylation of pRb (Figure 6D, S6I) and resulted in a transcriptional increase in the levels of E2F target genes (Figure 6E), suggesting a role for caspase-4 in cell cycle regulation by controlling p16^INK4a^ and p15^INK4b^ expression in OIS. Of note, activation of caspase-4 by intracellular LPS resulted in hypophosphorylated pRb and increased levels of E2F target genes (Figure S6J-K). Finally, similarly to SASP regulation, *GSDMD* targeting did not alter the cell cycle arrest in OIS (Figure S6L), suggesting that gasdermin-D has no significant role in OIS.

**Figure 6.**
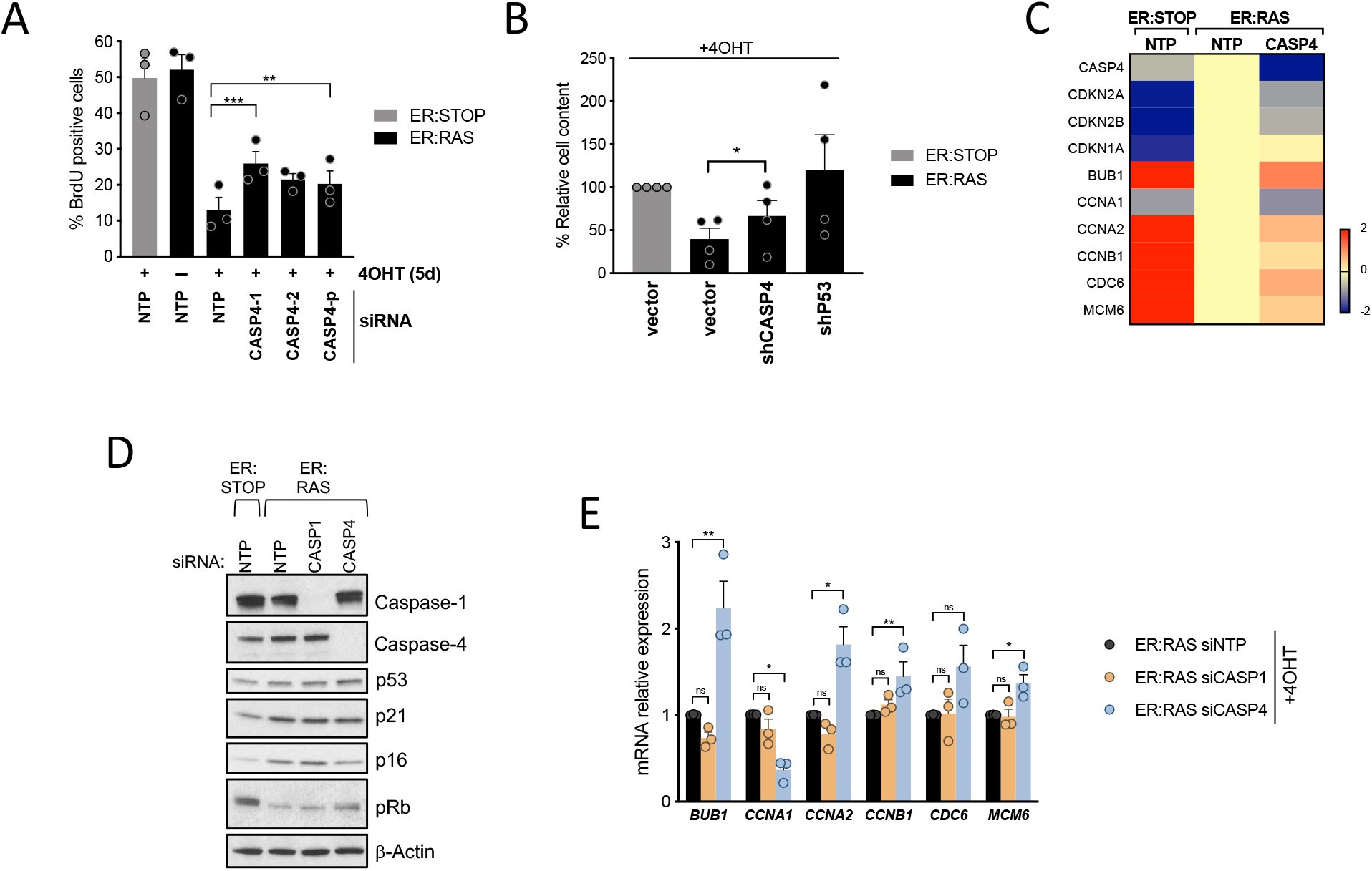
Caspase-4 contributes to the cell cycle arrest program in OIS. (A) IMR90 ER:STOP and ER:RAS cells were transfected with control (NTP), two individual *CASP4*-targeting siRNAs or a pool of 4 different siRNA sequences targeting *CASP4*, and treated with 4OHT or not as indicated. BrdU incorporation was measured by IF 5 days after 4OHT addition. (B) IMR90 ER:STOP/ER:RAS cells were stably transfected using retroviral shRNA vectors targeting *CASP4* or *TP53*. Infection with the empty vector (vector) was used as control. On day 0, equal number of cells were subjected to 4OHT treatment. Fifteen days after 4OHT addition, plates were fixed and stained with crystal violet. Crystal violet was extracted and used to quantify cell content. (C) Related to Figure 5A. DEG analysis between control ER:STOP and *CASP4*-knockdown ER:RAS compared to control ER:RAS after 5 days of 4OHT treatment was performed. Heatmap of the log2FC values from the indicated genes. (D) IMR90 ER:STOP and ER:RAS cells were transfected with control (NTP), *CASP1* or *CASP4*-targeting siRNAs and treated with 4OHT during 4 days. Cell lysates were subjected to immunoblotting analyses with the indicated antibodies. (E) mRNA relative expression of the indicated genes in IMR90 ER:RAS cells transfected with control (NTP) *CASP1* or *CASP4*-targeting siRNA was quantified by RT-qPCR after 5 days of 4OHT treatment. Statistical significance in A and B was calculated using two-tailed Student’s *t*-test. Statistical significance in E was calculated using one-way analysis of variance (ANOVA). \*\*\**P* < 0.001, \*\**P* < 0.01, \**P* < 0.05 and ns, not significant. All error bars represent mean ± s.e.m of 3 independent experiments.

Altogether, these results suggest that caspase-4 contributes to the proliferation arrest during senescence, impacting ultimately on the phosphorylation state of pRb resulting in transcriptional repression of E2F target genes.

### Caspase-11 is induced during cellular senescence in vivo

We have shown that caspase-4 expression levels are critical in cellular senescence. To investigate non-canonical inflammasome expression in senescence *in vivo*, we used three well-characterized mouse models of senescence. We first analysed the caspase-4 murine homologous caspase-11 expression in a model of OIS in which conditional expression of *Kras*^*G12D*^ by *Pdx-CRE* induces Pancreatic Intraepithelial Neoplasia (PanIN) in the pancreas of mice (Morton et al., 2010) (Figure 7A). We observed that low grade PanINs stained positive for caspase-11 when compared to surrounding pancreatic acinar cells, higher grade PanINs, and in the ducts and acinar cells in wild-type mice (Figure 7A). Importantly, quantification of Ki-67 staining in PanINs showed that the expression of caspase-11 was restricted to early senescent lesions with low proliferative index (Figure 7B), indicating that caspase-11 expression correlates with senescence in low-grade PanIN lesions.

**Figure 7.**
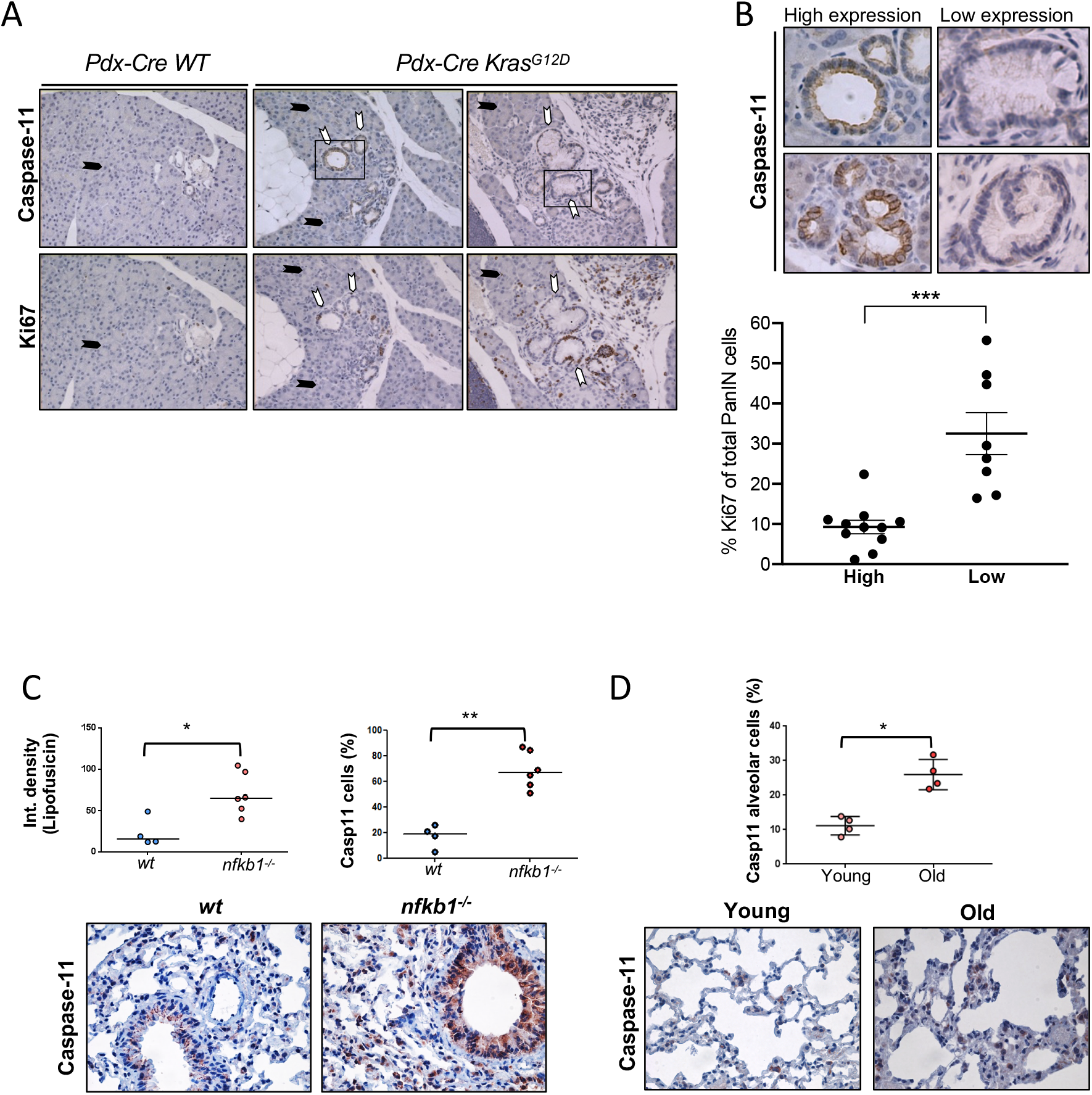
Caspase-11 expression is induced in senescence *in vivo*. (A) Immunohistochemistry showing Ki-67 and caspase-11 staining in sections from *Pdx-cre WT* and *Pdx-cre Kras*^*G12D*^ pancreas (left panels). Black arrows indicate acinar pancreatic cells, white arrows indicate PanIN cells. Close up images showing PanINs with high and low expression of caspase-11 (Right panel). (B) Quantification of Ki-67 positive cells of total PanIN cells from a total of 11 mice of 7 to 15 weeks of age. PanINs were classified according to the expression of caspase-11 as indicated. The percentage of Ki-67 positive cells was calculated scoring all cells of PanINs classified as high or low caspase-11 expression per mouse as indicated. Scatter plots were generated from total cells from high and low caspase-11 expressing PanINs with individual points representing the mean Ki-67 percentage positivity for each mouse, with horizontal lines representing group mean and s.e.m. Statistics: Mann Whitney U test. ***p < 0.001 (C) Analysis of caspase-11 expression was conducted by immunohistochemistry in lung sections from wild type (WT) or *nfkb1* knock out mice (*nfkb1*^*−/−*^) at 9.5 months of age. 10-15 random images were captured per mouse and average percentage positivity calculated for airway epithelial compartments. Scatter plots represent mean percentage positivity for each animal with horizontal line representing group median. Broad-band autofluorescence (an indicator of lipofuscin accumulation) was acquired from paraffin-embedded sections excited at 458 nm with fluorescence emission captured above 475 nm using a fluorescence microscope (Leica DM550B). Fluorescence intensity was analyzed using ImageJ. At least 10 small airways were analyzed per mouse and an average intensity calculated per animal. Scatter plots represent average value per animal with the horizontal line representing group median. Statistics: Mann Whitney U test. *p < 0.05, **p < 0.01 Representative images Casp4 staining by immunohistochemistry in airway epithelial cells from wt and *nfkb1−/−* mice, captured using x40 objective. (D) Analysis of caspase-11 expression by immunohistochemistry in lung sections of wt mice at 6.5 months of age (Young) and 24 months of age week (Old). Scatter plots were generated from 10-15 random images captured per animal with individual points representing mean percentage positivity for each mouse with horizontal line representing group median. Statistics: Mann-Whitney U test. *p < 0.05. Representative images of caspase-11 staining by immunohistochemistry (positive, brown; negative, blue) in alveolar cells from wt mice 6.5 and 24 months of age, captured using x40 objective.

We then investigated the expression of caspase-11 in two additional models of senescence *in vivo*. We detected increased expression of caspase-11 and the markers of senescence lipofuscin in lung airways in a model of inflammatory mediated activation of senescence by constitutive activation of NF-κB by knockout of its regulator *nfkb1*^*−/−*^ (*p50*^*−/−*^) (Jurk et al., 2014) (Figure 7C). Moreover, we observed an increase in the number of cells positively expressing caspase-11 in alveolar cells of the lung during organismal ageing (Figure 7D). In summary, these results support the model that non-canonical inflammasomes contribute to senescence *in vivo*.

## Discussion

Here, we show that cytoplasmic LPS recognition by the caspase-4 non-canonical inflammasome induces a senescence response, in which sublethal levels of LPS activate the p16^INK4a^-pRb and p53-p21^CIP1^ tumour suppressor pathways. Interestingly, while LPS-mediated pyroptosis requires caspase-4 and the effector protein gasdermin-D but not p53, the senescence response requires the participation of p53. These results suggest a mechanism in which p53 controls the cellular stress responses to microbial infection downstream of caspase-4 until a certain threshold level in which gasdermin-D-dependent pyroptosis eliminates highly damaged cells, introducing a new context-dependent role for p53 in innate immune sensing (Kastenhuber and Lowe, 2017). Further research will be necessary to determine the functional role of senescence in response to infection.

Our results describe a new mechanism of senescence induction by cytoplasmic microbial sensing, adding to the essential triggers such as the DNA Damage Response, telomere attrition, oncogenic activation, mitochondrial damage or ribosome biogenesis inactivation (Gorgoulis et al., 2019; Pantazi et al., 2019). We also show that both the expression and the assembly of the caspase-4 non-canonical inflammasome are triggered upon oncogene activation. Furthermore, we observe that caspase-4 is critical for SASP induction and contributes to the cell cycle arrest in OIS. Thus, these results suggest that the role of caspase-4 in senescence is conserved in a sterile context and that there is crosstalk between anti-microbial immune responses and tumour suppression. Notably, the assembly of the caspase-4 inflammasome is an early event in OIS, peaking after day three to four after Ras activation (Figure 4). Recent results suggest that OIS is a highly dynamic process, with distinct signalling waves contributing to the establishment of the senescent phenotype. Interestingly, the specific time point where caspase-4 is assembled coincides with the moment of transition to a proinflammatory, NF-κB - dependent SASP in OIS (Hoare et al., 2016) (Martinez-Zamudio et al., 2020). It is plausible that the activation of the non-canonical inflammasome could play a critical role in this transition.

Unexpectedly, our results indicate that LPS-mediated caspase-4 induced senescence is not accompanied by robust activation of IL-1β and the SASP. Interestingly, a significant SASP induction is only achieved when LPS stimulation and priming of the inflammasome with TLR2 ligands happen simultaneously, suggesting that the control of cellular senescence by caspase-4 is independent of the SASP. However, our data shows that caspase-4 heavily influences the SASP induction during OIS, where sustained production of A-SAA signaling through TLR2 is critical to the SASP (Hari et al., 2019). Thus, caspase-4 appears to be central regulating cell fate decisions occurring upon microbial-cytoplasmic recognition or sterile cellular stresses controlling the induction of cellular senescence, the SASP, or pyroptosis. Further research will be required to identify the nature of the signal responsible for caspase-4 activation during OIS.

Mechanistically, our data suggest that the role of caspase-4 in senescence is independent from its catalytic activity. Caspase-4 is a pattern recognition receptor that binds LPS directly (Shi et al., 2014); therefore, it is plausible that upon ligand recognition, caspase-4 functions as a scaffold platform for the activation of downstream senescence pathways in a proteolytic independent fashion. Our data suggests that pyroptosis is triggered only at a certain threshold of caspase-4 induction, suggesting a dose-dependent functional split between the caspase-4 pro-senescent and pyroptotic functions.

Inflammasomes have shown diverse pro- and anti-tumorigenic functions in cancer (Karki and Kanneganti, 2019). Here, we show that caspase-4 expression is induced following tumour initiation in a genetically engineered mouse model of pancreatic cancer and during ageing, suggesting a suppressive role for this pathway in cancer initiation. In recent years several strategies have been implemented to eliminate senescent cells or to modulate the activation of the SASP in anti-ageing and cancer therapies. (Baker et al., 2016; Baker et al., 2011; Dorr et al., 2013). Furthermore, the pharmacological targeting and removal of senescent cells has been shown to improve homoeostasis following tissue damage and ageing (Baar et al., 2017; Chang et al., 2016). Here we propose that manipulation of non-canonical inflammasomes could provide a new rationale for senotherapies and the implementation of pyroptosis for senolysis in cancer and ageing.

## Acknowledgements

We especially thank Maria Christophorou, Alex von Kriegsheim, Noor Gammoh, Andrew Finch, Simon Wilkinson, Manuel Collado, Claudia Wellbrock, Nick Hastie, Javier Caceres, Liz Patton and all the members of J.C.A. lab for helpful criticism, discussion and encouragement. This work was supported by Cancer Research UK (C47559/A16243 Training & Career Development Board - Career Development Fellowship), University of Edinburgh (Chancellor’s Fellowship) and the Wellcome Trust-ISSF. P.H., I.F.D and N.T. were funded by the University of Edinburgh Chancellor’s Fellowship. J.F.P and J.B. were funded by BBSRC (grant BB/K017314/1). F.R.M is funded by a Wellcome Trust Clinical Research Fellowship through the Edinburgh Clinical Academic Track (ECAT) (203913/Z/16/Z).

## Author contributions

Conceptualisation, I.F.D and J.C.A.; Formal Analysis, I.F.D, P.H, N.T., J.B., J.F.P. and J.C.A; Investigation, I.F.D, P.H., F.R.M., N.T., A.Q., J.B., and J.C.A.; Resources, J.C.A and V.G.B.; Data Curation, I.F.D. and J.C.A; Writing-Original Draft I.F.D and J.C.A.; Writing-Review & Editing, F.R.M.; Visualisation, I.F.D., N.T., J.B., J.F.P. and J.C.A; Supervision, J.C.A.; Funding Acquisition, J.C.A.

## Methods

### RESOURCE AVAILABILITY

#### Lead Contact and Materials Availability

Further information and requests for resources and reagents should be directed to and will be fulfilled by the Lead Contact, Juan-Carlos Acosta.

#### Data and Code Availability

The transcriptomic data generated during this study is available at GEO.

### EXPERIMENTAL MODEL AND SUBJECT DETAILS

HEK293T and IMR90 female human fetal lung fibroblast cells were obtained from American Type Culture Collection. All cell lines were maintained in Dulbecco’s Modified Eagle’s Medium (DMEM) (Sigma), supplemented with 10% Fetal Bovine Serum (FBS) (ThermoFisher) and 1% antibiotic-antimycotic solution (ThermoFisher). All cell lines were cultured at 37°C with 5% CO_2_ and tested for mycoplasma on a regular basis. All cell lines were regularly tested for mycoplasma contamination using the Mycoalert Mycoplasma Detection Kit (Lonza). Cell counting and viability were performed Muse® Count & Viability Assay Kit in a Muse Cell Analyser (Merck Millipore).

### METHOD DETAILS

#### Chemical compounds and treatments

OIS was induced by treating IMR90 ER:RAS cells with 100 nM 4OHT. IMR90 ER:RAS and control ER:STOP were maintained in standard media supplemented with 200 μg/ml geneticin. To induce oncogene induced senescence senescence, IMR90-ER:RAS cells were treated with 100 μM etoposide for 48 hours, followed by 5 days in normal culture media, For DNA damage-induced senescence, IMR90 cells were treated with 10 μM Etoposide for 48 hours. For non-canonical inflammasome activation and inflammasome priming experiments, ultrapure lipopolysaccharide (LPS) from E. coli 111:B4 (Invivogen), muramyldipeptide (MDP) (Tocris), Pam2CSK4 (Tocris), Pam3CSK4 (Tocris), Recombinant Human Apo-SAA (A-SAA)(Peprotech) and BSA (Sigma) were used. For priming time-course experiments, the following concentrations were used: LPS (1 μg/ml), MDP (1 μg/ml), Pam2CSK4 (50 ng/ml), Pam3CSK4 (500 ng/ml). To prime inflammasomes prior to LPS transfection, cells were treated with Pam2CSK4 (1 μg/ml) A-SAA (10 μg/ml) or BSA (10 μg/ml) for 3 hours.

#### Cell quantification and viability

To determine viable cell concentration of cultures, cells were washed and incubated with trypsin (ThermoFisher) for 5 min at 37 °C. Fully detached cells were collected by centrifugation, resuspended in Muse Count & Viability Reagent (Merck Millipore), and counted using the Muse Cell Analyzer (Merck Millipore). To determine cell viability, culture supernatants and attached cells were pooled together before centrifugation. Pellets were resuspended in Muse Count & Viability Reagent (Merck Millipore) and analysed using the Muse Cell Analyzer (Merck Millipore).

#### LPS transfection

To electroporate LPS or MDP, the Neon Transfection System (Invitrogen, MPK5000) and the Neon Transfection System 100 μL Kit (MPK10025) were used. Per each tip, 5 × 10^5 cells were transfected with the indicated amount of LPS or MDP. Electroporation parameters were set at 1500V, 30 ms and pulse number 1 for IMR90 cells and 1100V, 20 ms and pulse number 2 for HEK293T cells. Once electroporated, the tip content was unloaded into a clean Eppendorf tube and tubes were centrifuged on a bench-top centrifuge at 3000 rpm 3 min. Supernatant was removed to avoid any traces of MDP or LPS in the extracellular media prior to plating.

#### Conditioned medium for paracrine senescence transmission

For the production of conditional medium (CM), IMR90 ER:STOP and ER:RAS cells were cultured as previously described (Boumendil et al., 2019). IMR90 ER:STOP and ER:RAS cells were cultured with DMEM supplemented with 100 nM 4OHT and 10% FBS for 4 days, and with DMEM 100 nM 4OHT and 1% FBS for 4 additional days. CM was filtered using 0.2 μm syringe filters (Millipore) and reconstituted with a solution of DMEM supplemented with 40% FBS at a 3:1 ratio.

#### Generation of plasmids

Using standard retro-transcription procedures, total RNA extracted from IMR90 cells was converted into cDNA generating a human coding sequence (CDS) library. CASP1 and CASP4 CDS were amplified from the obtained library and cloned into the pMSCV-puro vectors The caspase-4 catalytically dead C258A pMSCV-puro vector was generated from the wild-type pMSCV-puro-CASP4 through site-directed mutagenesis by PCR using the Q5 Site-Directed Mutagenesis Kit (New England Biolabs). The CASP4 and GSDMD-targeting lentiviral vectors (pGIPZ) were purchased from Dharmacon. The CASP4-targeting retroviral vector (pRS-shCASP4) was generated inserting the oligonucleotide (+) GATCCCCCAACGTATGGCAGGACAAATTCAAGAGATTTGTCCTGCCATACGTTGTTTTTG into the pRS empty backgone following the pSuper RNAi System manual (OligoEngine) instructions. pLN-ER:RAS, LSXN-ER:Stop, MSCV-Ras^G12V^, pCMV-VSVG, and pUMVC3-gag-pol vectors have been described elsewhere (Acosta et al., 2013).

#### Retroviral and lentiviral production and infection

For retroviral production, 20 μg retroviral plasmid were cotransfected with 2.5 μg pCMV-VSVG envelope plasmid and 7.5 μg pUMVC3-gag-pol helper vector using polyethylenimine linear (Alfa Aesar) into HEK293T cells. For lentiviral production, 10 μg lentiviral plasmid were cotransfected with 2.5 μg pCMV-VSVG and 7.5 μg psPAX2 using polyethylenimine linear (Alfa Aesar) into HEK293T cells. Viral supernatant was collected from the HEK293T cells 2 days after transfection and passed through a 0.45 μm syringe filter (ThermoFisher). The viral supernantant was complemented with hexadimethrine bromide (Sigma) to a final concentration of 4 μg/mL. When performing retroviral infections, IMR90s were treated with fresh viral supernatant and subsequent viral supernatant collection and incubation of IMR90 cells was performed every 3 hours until three rounds of infection were performed. For lentiviral infections, a single 3 hour incubation with 1:10 dilution of viral supernatant was performed. In both cases viral supernatant was removed after the indicated rounds of infection, fresh media was added to IMR90 cells and, 2 days later, selection with puromycin (1 μg/ml) (ThermoFisher) was initiated. Before set-up, fresh standard media supplemented with the selection agent was added for over a week or until no alive cells were observed in control cells infected with a non-containing selection marker vector.

#### siRNA transfection

ON-TARGETplus siRNAs were obtained from Dharmacon. Sequences and IDs are detailed in the Key Resources Table. For all transfections, 30 nM siRNA were incubated up to 1 hour with Dharmafect 1 (Dharmacon, 1 μg/ml final use concentration) to allow the formation of siRNA:transfection agent complexes prior to transfection. On day 0, 200.000 IMR90 ER:STOP and ER:RAS cells were plated in each T-6 well, 4OHT was added and siRNA reverse transfections performed. Due to the transient nature of siRNA, cells were split 1:4 on day 3 and reverse transfections were repeated. To maintain the knockdown during 8 days, forward transfections were performed again on day 5.

#### Total RNA preparation and quantitative reverse transcription polymerase chain reaction (qRT-PCR)

Cell lysates were homogenized using QIAshredder (Qiagen) and RNA was extracted using the RNeasy Plus Mini kit (Qiagen). RNA was transformed into cDNA using qScript cDNA Supermix (Quanta Biosciences) following manufacturer’s instructions. To perform quantitative PCRs, samples were prepared in triplicates in 96-well plates. Each well contained 1 μL of cDNA, 200 nM forward primer, 200 nM reverse primer, 1x SYBR Select Master Mix (Applied Biosystems) and up to 20 μL of ultrapure DNase/RNase-free distilled water (ThermoFisher). Plates were loaded into a StepOnePlus Real-Time PCR System (ThermoFisher) and the following PCR cycling parameters were used: 10 min at 95 °C; 40 cycles of 15 s at 95 °C, 30 s at 60 °C and 15 s at 72 °C; 15 s at 95 °C. Data was analyzed using the double Delta Ct method. The housekeeping gene ACTB was used to normalize data. Primers are specified in the Table S1.

#### Immunofluorescence and high-content microscopy

Cells were fixed with 4% paraformaldehyde (FD NeuroTech) in PBS during 45 min. All incubations were performed at room temperature and on an orbital shaker. To permeabilize cells, cells were incubated with 0.2% Triton-X100 in PBS for 10 min. Cells were blocked with immunofluorescence blocking buffer (1% Bovine Serum Albumin (BSA) and 0.2% Fish Gelatin in PBS). Primary and secondary antibodies were diluted in immunofluorescence blocking. Anti-BrdU primary solution was supplemented with 0.5 U / μL DNAse (Sigma) and 1 mM MgCl2 to improve anti-BrdU access to DNA-bound BrdU. Nuclei were stained with 1 μg/mL 4’,6-diamidino-2-phenylindole (DAPI) (Molecular Probes). Antibodies are listed in the Key Resources Table. Immunofluorescence was analyzed using the high-contess High-Content Image Acquisition and Analysis software (Molecular Devices) as previously described (Boumendil et al., 2019). One-wavelength images of the same frame were merged using the software Fiji (ImageJ).

#### Western blot analysis

Whole cells were lysed in 1X Cell Lysis Buffer Cell (Cell Signalling) supplemented with cOmplete EDTA-free Protease Inhibitor Cocktail (Roche). Protein concentration was determined by the Bradford assay using the Bradford reagent (Biorad) and BSA pre-set standards (ThermoFisher) to construct a standard curve. To prepare samples for sodium dodecyl sulfate polyacrylamide gel electrophoresis (SDS-PAGE), 15 μg of protein were mixed with 6 μL 6x reducing Laemmli SDS sample buffer (Alfa Aesar) in a final volume of 36 μL. Samples were boiled 5 min at 95 °C and loaded in pre-cast Novex Tris-Glycine gels (Invitrogen). Pre-cast gels were run in an XCell SureLock™ Mini-Cell Electrophoresis tank (ThermoFisher) at 100 – 140 V. Proteins were transferred into nitrocellulose membranes using the iBlot Gel Transfer Device (ThermoFisher). Membranes were blocked in Tris-buffered saline (TBS) buffer (25 mM Tris-HCl + 137 mM NaCl + 2.7 mM KCl, pH7.4) supplemented with 5% non-fat milk 1 h at room temperature on a rocking shaker. Primary and secondary antibodies (described in the Key Resources Table) were diluted in TBS 5% milk buffer. To visualize bands, membranes were incubated with enhanced chemiluminescence solution (GE Healthcare) and exposed to X-ray films (GE Healthcare).

#### Caspase-4 fluorometric activity assay

To measure LEVD-AFC cleavage, the Caspase 4 Fluorometric Assay kit (Fluorometric) was used following manufacturer’s instructions. 2 × 10^6 IMR90 ER:STOP or ER:RAS cells were lysed in 50 μL cell lysis buffer. The assay was conducted in black sterile 96-well polystyrene plate (ThermoFisher). Fluorescence was measured (excitation filter: 400 nm; emission filter: 505 nm) using an Infinite^®^ 200 PRO (Tecan) plate reader.

#### Detection of caspase-4 oligomerization

Fresh IMR90 ER:STOP or ER:RAS cell pellets were resuspended in 0.5 ml of ice-cold buffer A (20 mM HEPES-KOH, pH 7.5; 10 mM KCl; 1.5 mM MgCl2; 1 mM EDTA; 1 mM EGTA; 320 mM sucrose), lysed by shearing 10 times through a 25-gauge needle, and centrifuged 8 min at 1.800 g at 4 °C. At this point, 30 μL of lysates were kept as input controls. Remaining supernatants were diluted with 1 volume of CHAPS buffer (20 mM HEPES-KOH, pH 7.5; 5 mM MgCl2; 0.5 mM EGTA; 0.1 mM PMSF; 0.1% CHAPS) and centrifuged 8 min at 5,000 x g. Supernatants were discarded and pellets were resuspended in 50 μL of CHAPS buffer containing 4 mM of disuccinimidyl suberate (DSS) during 30 min at room temperature to cross-ink proteins. Then, samples were centrifuged 8 min at 5,000 x g at 4 °C, supernatants discarded and pellets resuspended in 60 μL of protein loading buffer (25 mM Tris-HCl, pH 6.8; 1% SDS; 10% glycerol; 6.25 mM EDTA; 0.01% bromophenol blue). Samples were heated for 2 min at 90 °C and 18 μL of resuspended cross-linked pellets were loaded onto a 4-12% pre-cast Novex Tris-Glycine gels (Invitrogen). Further immunodetection of caspase-4 was performed following standard western blotting procedures.

#### Determination of IL-1β content in conditioned media

Conditioned media was collected, centrifuged 10 min at 1000 rpm at 4 °C and transferred to a clean tube. Released IL-1β was quantified using the Human IL-1 beta ELISA Ready-Set-Go! Kit (ThermoFisher) following the manufacturer’s instructions. Conditioned media IL-1β concentrations were deducted interpolating the data from the standard curve, as previously described (Fernandez-Duran et al., 2019).

#### Cell proliferation assays

5-bromo-2’-deoxyuridine (BrdU) incorporation was used to measure the number of cells actively replicating DNA. Cells were incubated with 10 μM BrdU (Sigma) for 16 to 18 hours. Cells were stained for immunofluorescence and high-content microscopy as described. To analyze long-term growth, low equal amounts of cells were plated in 10 cm diameter dishes. Media was changed every 3 days and cells were fixed two weeks after initial seeding with 0.5% glutaraldehyde (Sigma) in PBS for 20 min and left drying overnight. Dishes were stained with 0.2% crystal violet for 3 hours, washed twice with tap water and dried. To quantify cellular mass, cell-bound crystal violet was extracted in 10% acetic acid, equal amounts were transferred to a spectrophotometer-compatible 96-well plate and absorbance was read at 595 nm.

#### SA-β-Galactosidase assay

5 × 10^4^ IMR90 cells per well were seeded in 6-well plates. Four days later, cells were fixed with 0.5% glutaraldehyde (Sigma) in PBS during 10 min. Fixed cells were washed three times with PBS 1 mM MgCl2 pH 5.7, before adding to each well 2 mL of pre-warmed X-Gal staining solution (2 mM MgCl2, 5 mM K4Fe(CN)6 • 3H2O, 5 mM K3Fe(CN)6, 1 mg/mL X-Gal solution ready to use (ThermoFisher) in PBS). Plates were incubated for 2-24 h at 37 °C, washed and imaged. SA-β-Gal activity positive and negative cells were quantified using FIJI/ImageJ.

#### AmpliSeq transcriptome profiling

RNA quality was assessed on the Bioanalyser 2100 Electrophoresis Instrument (Agilent) with the RNA 6000 Nano Kit (Agilent). Samples were quantified using the Qubit 2.0 fluorometer and the Qubit RNA Broad Range assay.10 ng of RNA was reverse-transcribed to cDNA, and target genes were amplified for 12 cycles of PCR using the Ion AmpliSeq Human Gene Expression Core Panel (Thermofisher). This panel contains a pool of 20,802 amplicons (41,604 primers) of approximately 150 bases in length. Ion Torrent sequencing adapters and barcodes Ion XpressTM Barcode Adapters (Ion XpressTM Barcode Adapters) were ligated to the amplicons and adapter-ligated libraries were purified using AMPure XP beads. Libraries were quantified by qPCR and diluted to 100 pM before being combined in equimolar pools of 8 per each Ion PI Chip Kit v3 (ThermoScientfic). Sequencing was performed using the Ion PI Hi-Q Sequencing 200 Kit (ThermoFisher). Sequence reads were mapped to the hg19_AmpliSeq_Transcriptome_ERCC_v1.fasta reference. BAM files were generated using the Torrent Suite software v 5.2.0 (ThermoFisher). Differentially expressed gene (DEG) analysis was performed with the DESeq2 package v.1.20.0. Gene Set Enrichment Analysis (GSEA) was performed using the Broad Institute GSEA software v3.0. DEG-obtained log2FC values were used as inputs for the GSEA. Molecular signatures were obtained from MSigDB v.6.2.

#### Experiments with mice

Experiments were performed according to UK Home Office regulations. Mice carrying a conditional *Pdx1–Cre Kras*^*G12D/+*^ allele were used and have been described previously (Morton et al., 2010). Sections of formalin-fixed paraffin-embedded mouse pancreas from 6 to 14-week-old mice were stained with antibody against Ki67 and Caspase-4. Caspase-4 signal was used to classify high and low Caspase-4 expressing PanIN. Aging experiments were carried out on male wild-type C57BL/6 mice or male *nfkb1*^*−/−*^ mice on a pure C57BL/6 background at 6.5, 9.5 and 24 months of age.

#### Immunohistochemistry

For the Pdx1-Cre Kras^G12D/+^ mice (PanIN), Using EnVision™ +Dual Link system-HRP (DAB+) kit (K4065, Dako), sections of formalin-fixed paraffin-embedded mouse pancreas were stained with antibody against Ki67 (ab21700, Abcam). The total number of Ki67 positive cells per PanIN, and the total cells per PanIN were counted, and thus the percentage of Ki67 positive cells per PanIN was calculated. The mean score for each mouse was calculated and these scores were plotted scatter plot. Consecutive sections were stained with antibodies against Caspase 4/11 (bs-6858R, Bioss). The stainings were examined and classified for high or low expression of the respective antibodies, and each structure compared with the Ki67 percentage.

For nfkb1−/− and aging mice analysis, sections were dewaxed in histoclear (5 min), rehydrated through graded ethanol solutions (100, 90, and 70%) and washed in distilled H2O. Endogenous peroxidase activity was blocked by immersing sections in 0.3% H2O2 (Sigma, H1009) diluted in H2O for 30 min. To retrieve antigens, sections were boiled in 0.01 M citrate (pH 6.0). Sections were blocked in normal goat serum diluted 1:60 in 0.1% BSA in PBS. Sections were incubated with the primary antibody overnight at 4°C for Caspase 4/11 (bs-6858R, Bioss). Biotinylated secondary antibody was added and detected using the rabbit peroxidase ABC kit (Vector Laboratories, PK-4001), according to the manufacturer’s instructions. Substrate was developed using the NovaRed kit (Vector Laboratories, SK-4800).

Nuclei were counterstained with heamatoxylin, and sections were dehydrated through graded ethanol solutions, cleared in xylene, and mounted in di-nbutylehthalate in xylene (Thermo Scientific, LAMB-DPX). Staining was analysed with a NIKON ECLIPSE-E800 microscope, and images were captured with a Leica DFC420 camera using the LAS software (Leica). 10-15 random images were captured per section and the percentage of positively stained cells determined from total number of cells before an average per mouse was calculated.

#### Broad-band autofluorescence (lipofuscin accumulation) analysis

Broad-band autofluorescence was acquired from sections cut at 3 μm using X20 objective (Leica DM550B). Sections were excited at 458 nm and fluorescence emission captured above 475 nm. Fluorescence intensity per airway epithelium was quantified using ImageJ software and divided by background emission. At least 10 small airways per mouse was analysed.

### QUANTIFICATION AND STATISTICAL ANALYSIS

GraphPad Prism 7 software was used for statistical analysis. Results were displayed as the means ± SEM and statistical significance was determined with Student’s t tests, One-way analysis of variance (ANOVA) or Two-way ANOVA.

### KEY RESOURCES TABLE

| REAGENT or RESOURCE | SOURCE | IDENTIFIER |
| --- | --- | --- |
| **Antibodies** |  |  |
| Anti-Caspase-4 | Santa Cruz | 4B9 |
| Anti-BrdU | BD Biosciences | 555627 |
| Anti-IL-1α | R&D | MAB200 |
| Anti-IL-1β | R&D | MAB201 |
| Anti-IL-6 | R&D | AF206NA |
| Anti-IL-8 | R&D | MAB208 |
| Anti-p21 | Sigma | P1484 |
| Anti-p16 (IF) | ProteinTech | 10883-1-AP |
| Mouse-Alexa Fluor 488 | ThermoFisher | A-11029 |
| Rabbit-Alexa Fluor 594 | ThermoFisher | A-11037 |
| Goat-Alexa Fluor 594 | ThermoFisher | A-11058 |
| Anti-Caspase-1 | Adipogen | Bally-1 |
| Anti-Gasdermin-D | Novus Biologicals | NBP2-33422 |
| Anti-p16 (WB) | Santa Cruz | JC-8 |
| Anti-p53 | Santa Cruz | DO-1 |
| Anti-pRb | BD Pharmingen | 554136 |
| Anti-β-Actin | Santa Cruz | I-19 |
| Anti-Mouse-HRP | Sigma | A2554 |
| Anti-Rabbit-HRP | Sigma | A0545 |
| Anti-Goat-HRP | Sigma | 2020 |
| Anti-Mouse IgG (Fc specific)-Peroxidase | Sigma | Cat #A2554 |
| Anti-Rabbit IgG (whole molecule)-Peroxidase | Sigma | Cat #A0545 |
| Donkey anti-goat-HRP | Santa Cruz | Cat #sc-2020 |
| Goat anti-Rabbit IgG Alexa Fluor 594 | Thermo Fisher | Cat #A11037 |
| Monoclonal anti-B-Actin-peroxidase (AC-15) | Sigma | Cat #A3854 |
| Mouse monoclonal IgG2B Isotype Controls (20116) | R&D systems | Cat #MAB004 |
| Rabbit polyclonal anti-Caspase 4 + Caspase 11 Biotin conjugated | Bioss | Cat #bs-6858R |
| **Chemicals, Peptides, and Recombinant Proteins** |  |  |
| 4-hydroxytamoxifen | Sigma | H7904 |
| Geneticin | ThermoFisher | 10131-027 |
| Puromycin | ThermoFisher | A11138 |
| Ultrapure lipopolysaccharide (LPS) from E. coli 111:B4 | Invivogen | tlrl-3pelps |
| Muramyl dipeptide (MDP) | Tocris | tlrlmdp |
| Pam2CSK4 | Tocris | 4637 |
| Pam3CSK4 | Tocris | 4633 |
| Recombinant Human Apo-SAA | Peprtech | #300-13 |
| 5-bromo-2'-deoxyuridine (BrdU) | Sigma | 858811 |
| X-Gal solution ready to use | ThermoFisher | R0941 |
| **Critical Commercial Assays** |  |  |
| Neon Transfection System 100 μL Kit | Invitrogen | MPK10025 |
| Q5 site directed mutagenesis kit | New England Biolabs | E0554S |
| Caspase 4 Assay kit (Fluorometric) | Abcam | ab65658 |
| Human IL-1 beta ELISA Ready-Set-Go! Kit | ThermoFisher | 15581087 |
| RNA 6000 Nano Kit | Agilent | 5067-1511 |
| Ion AmpliSeq Human Gene Expression Core Panel | ThermoFisher | A26325 |
| Ion PI Hi-Q Sequencing 200 Kit | ThermoFisher | A26433 |
| **Deposited Data** |  |  |
| Ampliseq data | GEO |  |
| **Experimental Models: Cell Lines** |  |  |
| HEK293T | ATCC | N/A |
| IMR90 human fetal primary fibroblasts | ATCC | N/A |
| **Experimental Models: Organisms/Strains** |  |  |
| Mouse: C57BL/6 | The Jackson Lab. |  |
| **Oligonucleotides** |  |  |
| ON-TARGETplus Non-targeting Pool | Dharmacon | D-001810-10 |
| ON-TARGETplus CASP1-targeting Pool | Dharmacon | LQ-004401-00 |
| ON-TARGETplus CASP4-targeting Pool | Dharmacon | LQ-004404-00 |
| ON-TARGETplus CASP4-targeting #1 | Dharmacon | J-004404-06 |
| ON-TARGETplus CASP4-targeting #2 | Dharmacon | J-004404-08 |
| ON-TARGETplus GSDMD-targeting Pool | Dharmacon | LQ-016207-00 |
| The sequences and description of primers used for RT-qPCR are provided in Table S1 | N/A | N/A |
| **Recombinant DNA** |  |  |
| pMSCV-puro-CASP1 | This paper | Exp vector |
| pMSCV-puro-CASP4 | This paper | Exp vector |
| pMSCV-puro-CASP4-C258A | This paper | Exp vector |
| pGIPZ-shCASP1 | Dharmacon | V3LHS_392179 |
| pGIPZ-shCASP4 | Dharmacon | V3LHS_338745 |
| pGIPZ-shGSDMD | Dharmacon | V3LHS_378066 |
| pGIPZ shTP53 | Dharmacon | V3LHS_333920 |
| pRS-shCASP4 | This paper | N/A |
| pRS-shTP53 | (Acosta et al., 2008) | N/A |
| **Software and Algorithms** |  |  |
| MetaXpress High-Content Image Acquisition and Analysis software (used for quantification of immunofluorescence microscopy images) | Molecular Devices | N/A |
| FIJI/ImageJ (used for merging of immunofluorescence microscopy images and SA-β-Gal quantification) | ImageJ | N/A |
| Torrent Suite software v.5.2.0 | ThermoFisher | N/A |
| DESeq2 package v.1.20.0 | (Love et al., 2014) | Bioconductor |
| GSEA software v.3.0 | (Subramanian et al., 2005) | UC San Diego / Broad Institute |
| MSigDB v.6.2 | (Subramanian et al., 2005) | UC San Diego / Broad Institute |
| GraphPad Prism 7 | GraphPad Software | Statistics software |

## Supplemental information

**Figure S1.**
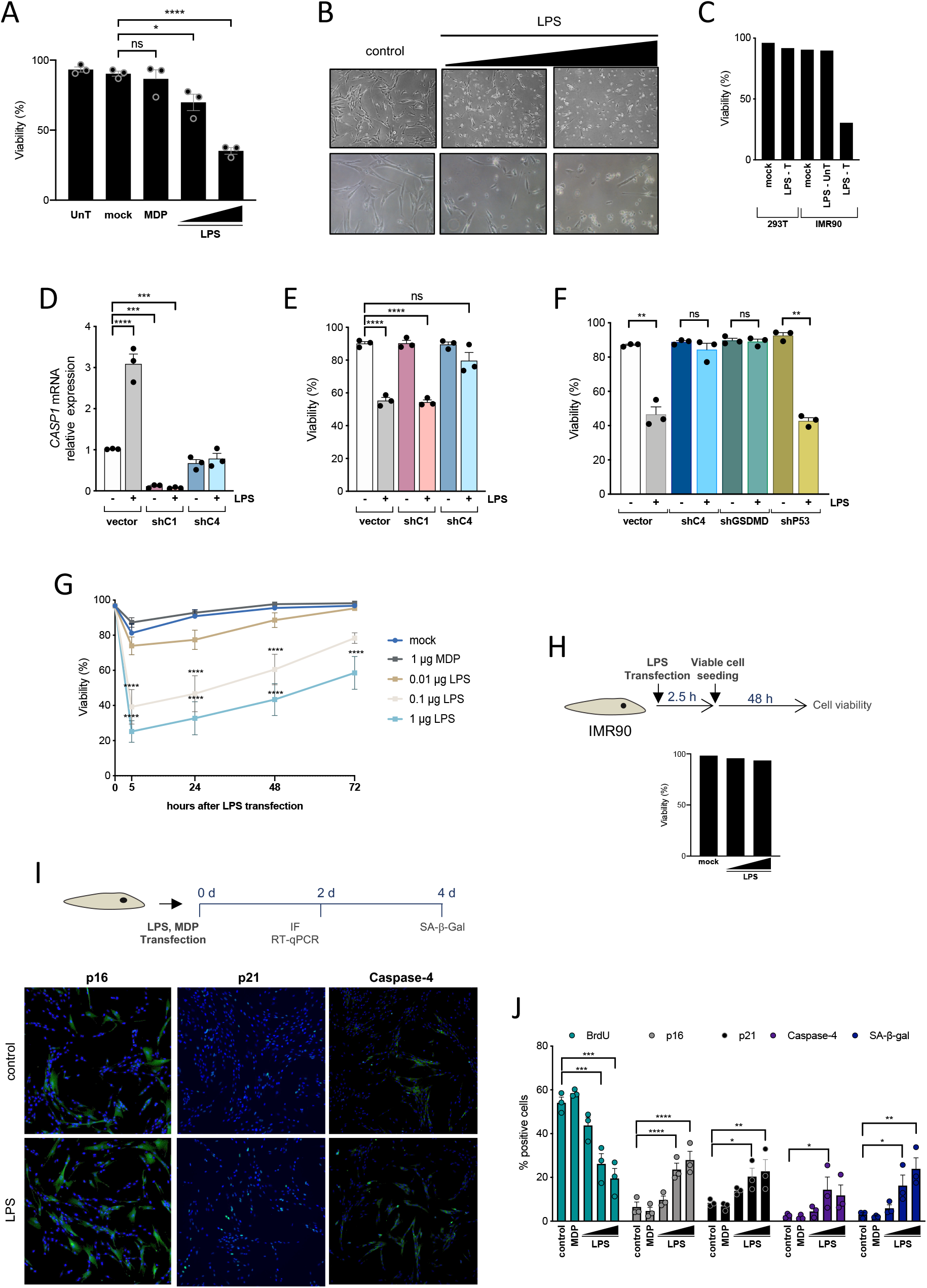
LPS-mediated caspase-4 activation induces a senescent phenotype in human primary fibroblasts – related to Figure 1. (A) IMR90 fibroblasts were un-transfected (UnT), mock-transfected (mock), transfected with MDP (1 μg / 5 × 10^5 cells) or increasing concentrations of LPS (0.1 or 1 μg LPS / 5 × 10^5 cells). Cell viability was measured 2 h after transfection. (B) Representative images of IMR90 cells mock-transfected (left) or transfected with 0.1 μg LPS / 5 × 10^^5^ cells (middle) or 1 μg LPS / 5 × 10^^5^ cells (right) under brightfield microscopy 24 h after transfection. (C) IMR90 or 293T cells were un-transfected (mock) or transfected with 1 μg LPS / 5 × 10^^5^ cells (LPS-T). To confirm that pyroptosis is dependent on intracellular localization of LPS, IMR90 were also treated with 1 μg LPS / 5 × 10^^5^ cells without further transfection (LPS - UnT). Cell viability was measured 2.5 h after transfection. Data from a single representative experiment. (D) Cells were treated as shown in Figure 1A. *CASP1* mRNA relative expression was quantified by RT-qPCR 48 h after LPS transfection. (E) Cells were treated as shown in Figure 1A. Cell viability was measured 24 h after LPS transfection. (F) IMR90 cells were stably infected with an empty pRS vector (vector) or a pRS vector targeting either *CASP4* (shC4), *GSDMD* (shGSDMD) or *TP53* (shP53) prior to transfection with 0.1 μg LPS / 5 × 10^^5^ cells, and cell viability was measured 24 h after LPS transfection. (G) IMR90 fibroblasts were mock-transfected, transfected with 1 μg MDP / 5 × 10^5 cells, or with increasing concentrations of LPS (0.01, 0.1 or 1 μg LPS / 5 × 10^5 cells). Cell viability was measured 5, 24, 48 and 72 h after transfection. (H) IMR90 fibroblasts were mock-transfected or transfected with increasing concentrations of LPS (0.1 or 1 μg LPS / 5 × 10^5 cells). Cell viability was measured 2.5 h after transfection and viable cells were replated and cultured for further 48 h before measuring cell viability again. Bars show a single representative experiment. (I-J) IMR90 fibroblasts were mock-transfected (control), transfected with 1 μg MDP / 5 × 10^5 cells, or with increasing concentrations of LPS (0.01, 0.1 or 1 μg LPS / 5 × 10^5 cells). BrdU incorporation, p21^CIP1^, p16^INK4a^, p21^CIP1^ and caspase-4 levels were measured by IF 48 h after transfection. SA-β-Gal activity was determined 4 days after transfection. Representative pictures (left) of IF staining for p16^INK4a^, p21^CIP1^ and caspase-4 of mock-transfected (control) and cells transfected with LPS (0.01 LPS / 5 × 10^5 cells) are shown. Statistical significance in A, D, E and J was calculated using one-way analysis of variance (ANOVA). Statistical significance in G was calculated using using two-way analysis of variance (ANOVA). Statistical significance in G was calculated using two-tailed Student’s *t*-test. ****P < 0.0001, \*\*\**P* < 0.001, \*\**P* < 0.01, and \**P* < 0.05. ns, not significant. All error bars represent mean ± s.e.m of 3 independent experiments.

**Figure S2.**
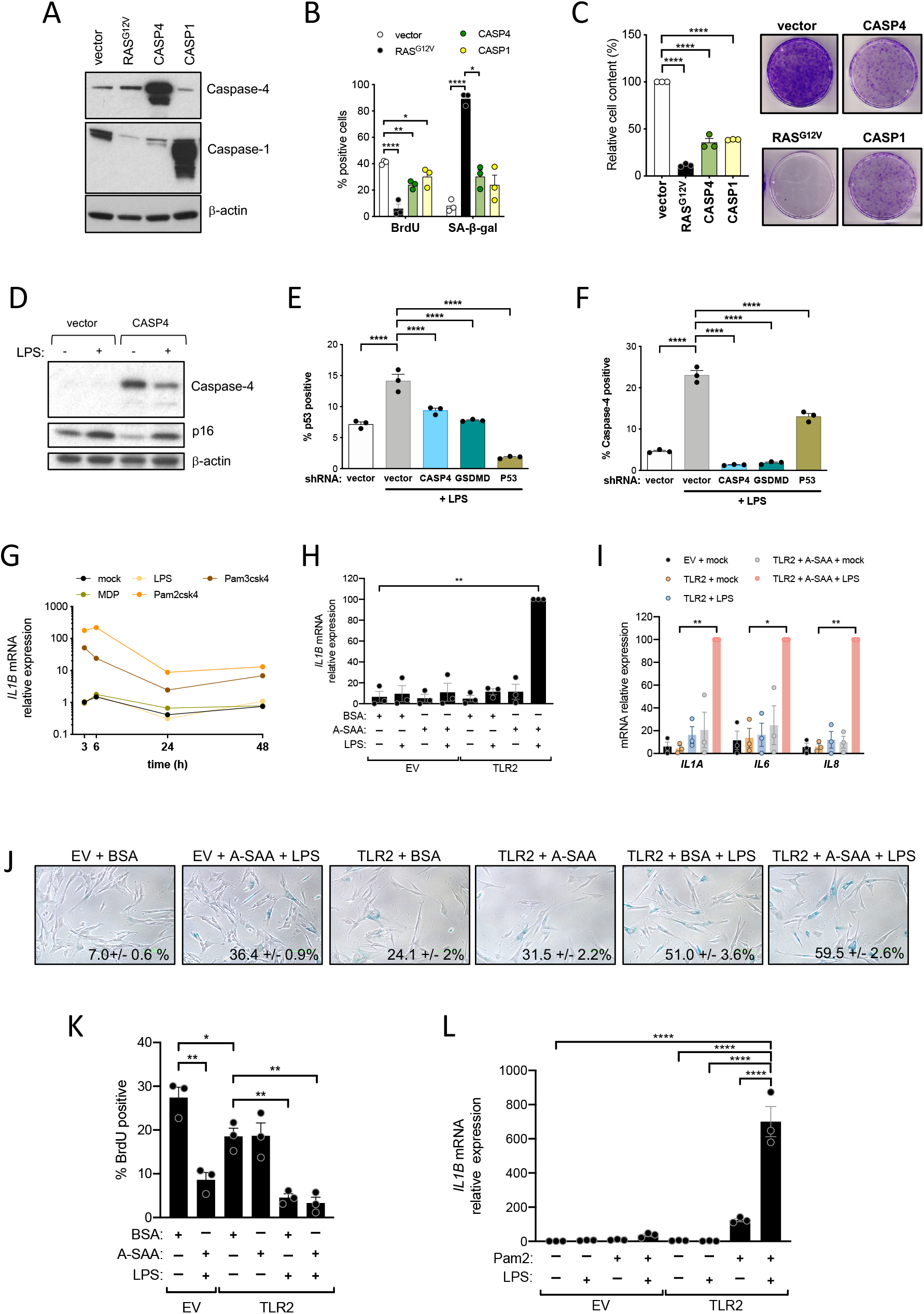
LPS-mediated caspase-4 induced senescence is independent on inflammasome priming – related to Figure 2. (A-C) IMR90 were infected with *CASP4*, *CASP1*, *RAS*^G12V^ expression vectors or empty vector (vector) control. (A) Protein amounts of CASP4 and CASP1 were analyzed by immunoblotting. (B) BrdU incorporation and SA-β-Gal activity were determined 4 days after seeding equal number of cells. (C) Relative cell content (left) was quantified 15 days after equal number of cells were seeded; representative images (right) of crystal violet stained cells are shown. (D) IMR90 cells were infected with *CASP4* expression vector or empty vector (vector) control and transfected with LPS (0.1 μg LPS / 5 × 10^5 cells). Caspase-4 and p16^INK4a^ protein levels were analyzed by immunoblotting 48 h after transfection. (E-F) Cells were treated as shown in Figure 1A. p53 (E) and caspase-4 (F) levels were measured by IF 48 h after LPS transfection. (G) IMR90 cells were treated with MDP (1 μg/mL), LPS, (1 μg/mL) Pam2CSk4 (0.05 μg/mL) and Pam3CSk4 (0.5 μg/mL). *IL1B* mRNA relative expression was quantified at the indicated points. Data from a single representative experiment. (H) IMR90 cells infected with TLR2 expressing vector or control empty vector (EV) were primed with 10 μg/ml of A-SAA for 3 hours prior to electroporation with LPS (1 μg/mL) to activate CASP4. *IL1B* mRNA relative expression was quantified 48 hours after LPS transfection. (I) Samples from (H) were analyted for *IL1A*, *IL6*, *IL8* mRNA relative expression48 hours after LPS transfection. (J) SA-β-GAL staining was conducted 48 hours after treatment as in (H). Values represent the Mean ± SEM of 3 independent experiments. (K) Proliferation capacity in experiment (H) was measured by BrdU incorporation assay. (L) Analysis of *IL1B* mRNA expression by qRT-PCR in IMR90 cells infected with TLR2 expressing vector or control empty vector (EV), primed with 1 μg/ml Pam2CSK4 for 3 hours, followed by electroporation with 1 μg/mL LPS for 48 hours. Statistical significance was calculated using one-way analysis of variance (ANOVA). ****P < 0.0001, \*\*\**P* < 0.001, \*\**P* < 0.01, and \**P* < 0.05. ns, not significant. All error bars represent mean ± s.e.m of 3 independent experiments.

**Figure S3.**
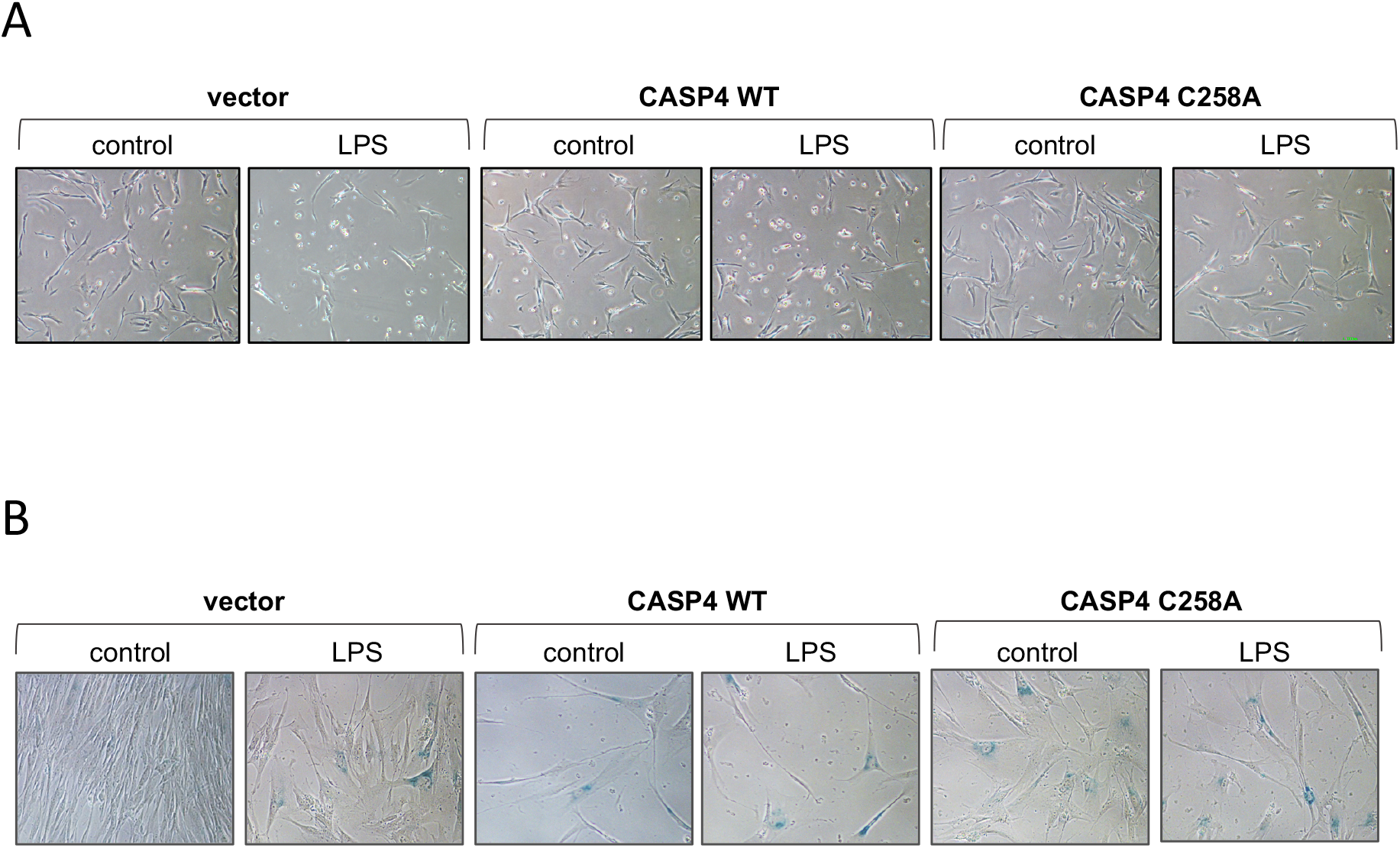
Caspase-4 mediated regulation of senescence is independent of its catalytical function – related to Figure 3. (A) Related to Figure 3D. Cells in culture under brightfield microscopy 24 h after transfection, at the time of viability assessment. (B) Related to Figure 3F. Representative images of SA-β-Gal stained cells 4 days after transfection.

**Figure S4.**
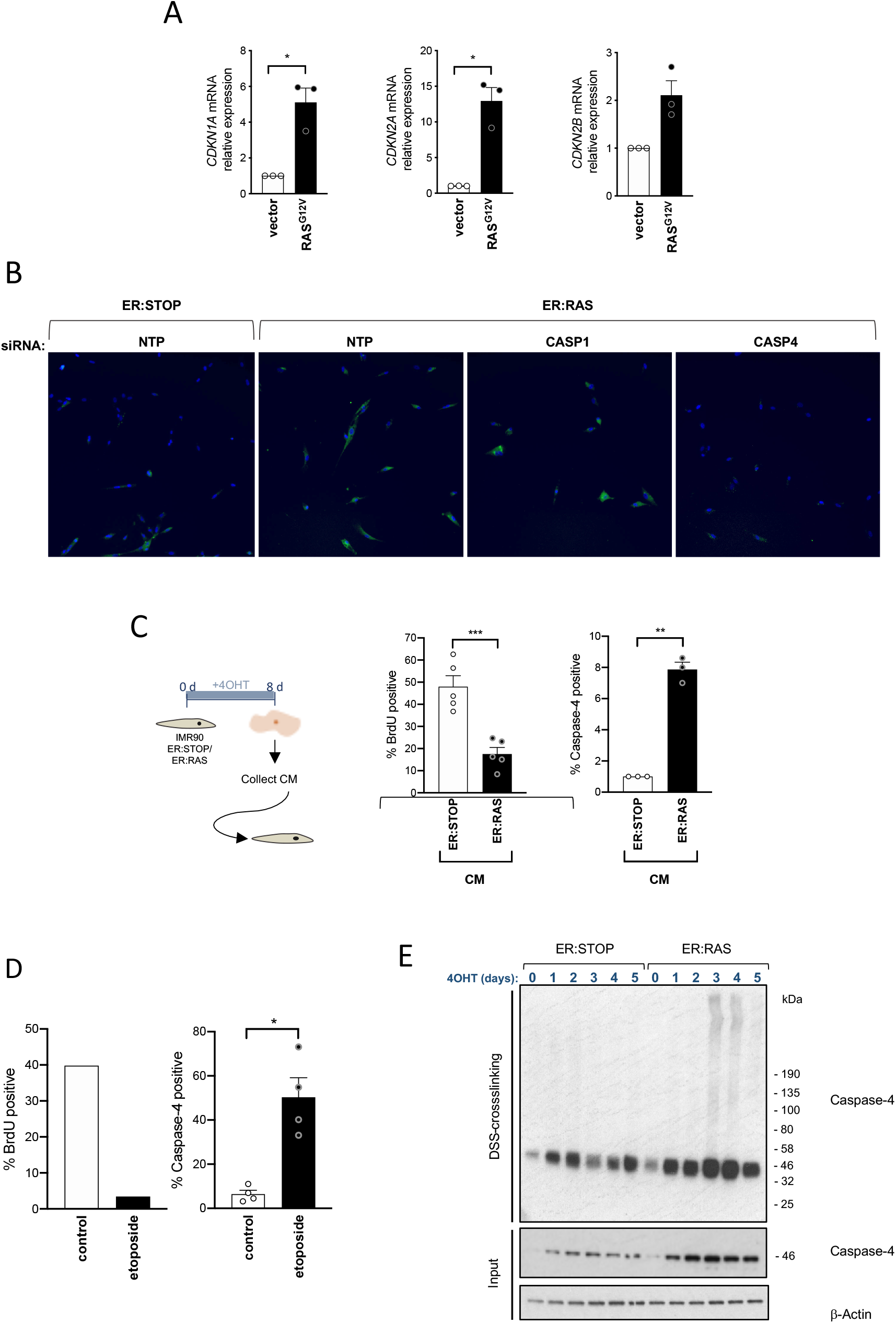
The caspase-4 non-canonical inflammasome is activated in Oncogene-induced senescence – related to Figure 4. (A) *CDKN1A* (p21^CIP1^), *CDKN2A* (p16^INK4a^) and *CDKN2B* (p15^INK4b^) relative expression levels were measured in IMR90 cells undergoing *RAS*^G12V^-OIS and control cells. (B) Representative images of caspase-4 stained cells by IF of IMR90 ER:STOP and ER:RAS cells treated with the indicated siRNA 5 days after the addition of 4OHT. (C) After 8 days of 4OHT treatment, conditioned media (CM) from IMR90 ER:STOP and ER:RAS cells was collected and added to IMR90. After 48 h, BrdU incorporation (middle) and caspase-4 levels (right) were measured by IF. (D) IMR90 cells were treated with 10 μM etoposide and 48 hour later BrdU incorporation (left) and caspase-4 (right) levels were measured by IF. Data relative to BrdU incorporation belongs to a single representative experiment. (E) IMR90 ER:STOP and ER:RAS cells were treated with 4OHT for the indicated time, cells were harvested and subjected to DSS-crosslinking. After SDS-PAGE separation, both DSS-crosslinked samples and inputs were probed for caspase-4 following western blot procedures. Statistical significance was calculated using two-tailed Student’s *t*-test. ****P < 0.0001, \*\*\**P* < 0.001, \*\**P* < 0.01, and \**P* < 0.05. ns, not significant. All error bars represent mean ± s.e.m of 3 independent experiments.

**Figure S5.**
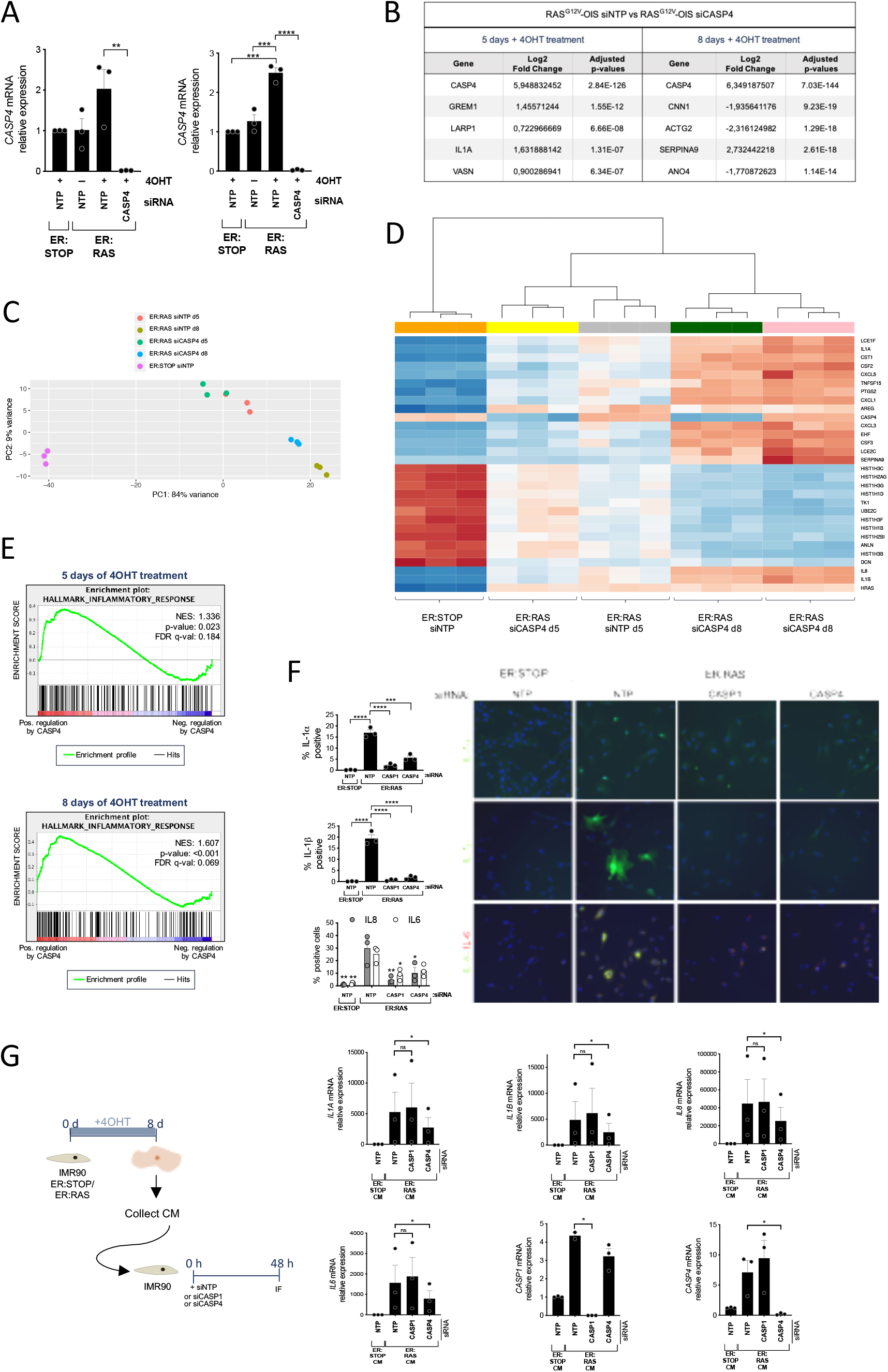
Caspase-4 activation controls the proinflammatory SASP – related to Figure 5. (A) *CASP4* mRNA relative expression levels were quantified by RT-qPCR after 5 days (left) and 8 days (right) of 4OHT treatment in ER:STOP and ER:RAS cells transfected with the indicated siRNA. (B) Top five differentially expressed genes upon *CASP4*-targeting in *RAS*^G12V^-OIS identified by DEG analysis 5 and 8 days after 4OHT treatment. (C) Principal component analysis (PCA) of variance stabilized transformed data using a parametric fit for the dispersion. Each dot corresponds to a sample replicate. (D) Heatmap and hierarchical clustering of the 30 genes with highest variance across all samples based on the total transformed data. (E) Related to Figure 5B. Enrichment plots of the signature “INFLAMMATORY RESPONSE” upon *CASP4*-targeting in RAS^G12V^-OIS 5 (top) and 8 (bottom) days after 4OHT treatment are shown. (F) IMR90 ER:STOP or ER:RAS cells were transfected with control (NTP), *CASP1* or *CASP4*-targeting siRNA and treated with 4OHT during 8 days. IL-1α, L-1β, IL-6 and IL-8 levels were analyzed by IF 8 days after the addition of 4OHT. Representative images as used for the high content analysis are also shown. (G) After 8 days of 4OHT treatment, conditioned media (CM) from IMR90 ER:STOP and ER:RAS cells was collected and added to IMR90. Concomitantly, IMR90 cells were transfected with control (NTP), *CASP1* or *CASP4*-targeting siRNA. After 48 h, *IL1A*, *IL1B*, *IL8*, *IL6*, *CASP1* and *CASP4* mRNA relative expression levels were measured by RT-qPCR. Statistical significance in A and F was calculated using one-way analysis of variance (ANOVA). Statistical significance in G was calculated using two-tailed Student’s *t*-test. ****P < 0.0001, \*\*\**P* < 0.001, \*\**P* < 0.01, \**P* < 0.05 and ns, not significant. All error bars represent mean ± s.e.m of 3 independent experiments.

**Figure S6.**
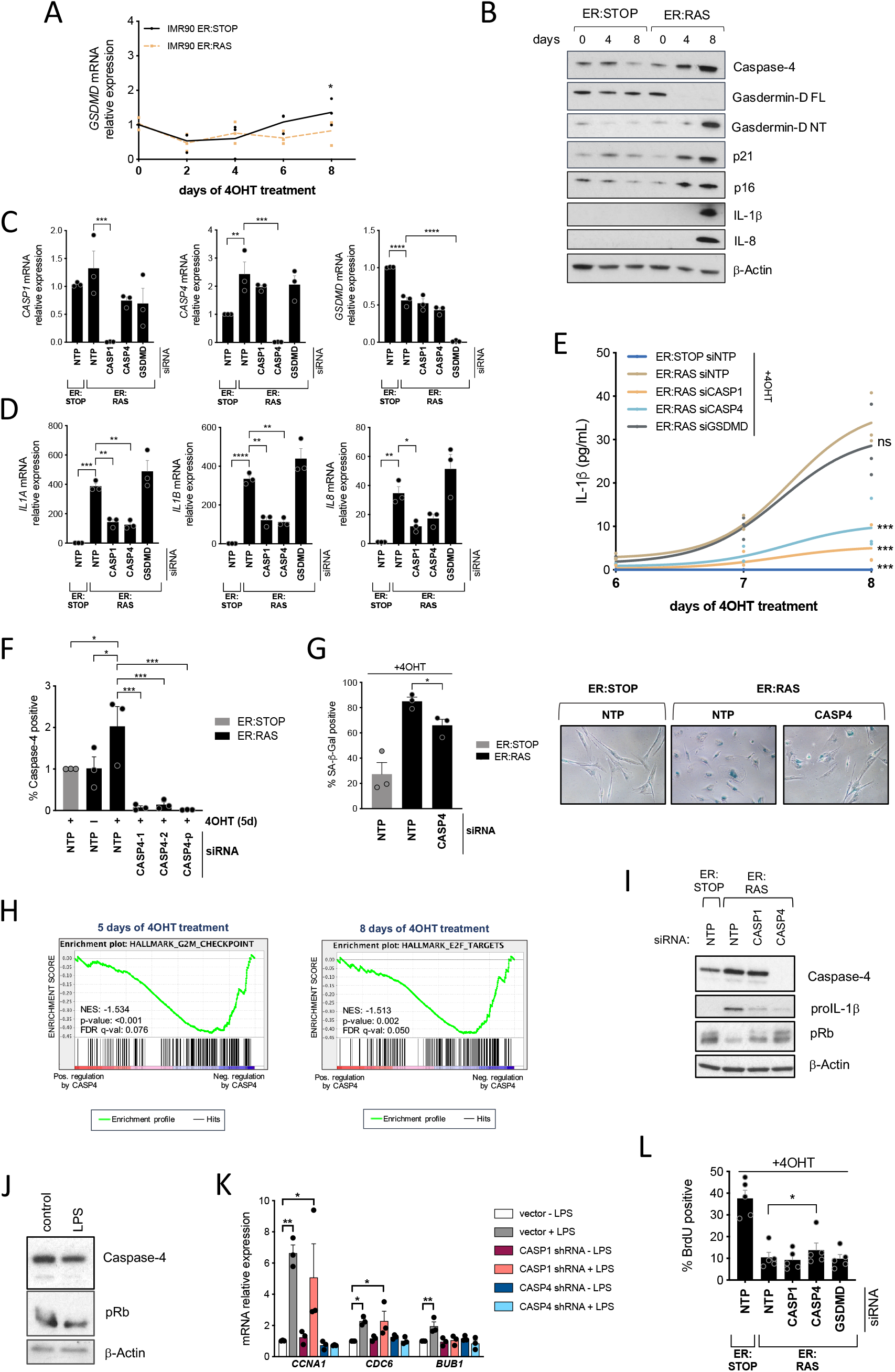
Caspase-4 contributes to the cell cycle arrest program in OIS – related to Figure 6. (A) *GSDMD* mRNA relative expression was quantified by RT-qPCR in IMR90 ER:STOP and ER:RAS cells 0, 2, 4, 6 and 8 days after 4OHT addition. (B) IMR90 ER:STOP/ER:RAS cells were treated with 4OHT for the indicated time. Caspase-4, full-length (FL) and N-terminal (NT) Gasdermin-D, p21^CIP1^, p16^INK4a^, IL-1β and IL-8 levels were analyzed by immunoblotting. (C) IMR90 ER:STOP/ER:RAS cells were transfected with control, *CASP1*, *CASP4* or *GSDMD*-targeting siRNAs. To analyze knockdown efficiency, *CASP1*, *CASP4* and *GSDMD* mRNA relative expression levels were quantified by RT-qPCR after 5 days of 4OHT treatment. (D) IMR90 ER:STOP and ER:RAS cells were transfected with control, *CASP1*, *CASP4* or *GSDMD*-targeting siRNAs. *IL1A*, *IL1B* and *IL8* mRNA relative expression levels were quantified by RT-qPCR after 5 days of 4OHT treatment. (E) IMR90 ER:STOP and ER:RAS cells were transfected with the indicated siRNAs and secreted IL-1β was quantified by ELISA 6,7, and 8 days after 4OHT addition. Statistical analysis was performed comparing control senescent cells (ER:RAS siNTP) to the other conditions 8 days after 4OHT addition. (F) IMR90 ER:STOP and ER:RAS cells were transfected with control (NTP), two individual *CASP4*-targeting siRNAs or a pool of 4 different siRNA sequences targeting *CASP4*, and treated with 4OHT or not as indicated. Caspase-4 levels were measured by IF 5 days after 4OHT addition. (G) IMR90 ER:STOP and ER:RAS cells were transfected with control, or *CASP4*-targeting siRNAs and SA-β-Gal activity was determined 8 days after the addition of 4OHT (left). Representative images for SA-β-Gal activity are shown (right). (H) Enrichment plots of the signatures “G2M CHECKPOINT” (left) and “E2F TARGETS” (right) upon *CASP4*-targeting in *RAS*^G12V^-OIS IMR90 cells 5 days after 4OHT treatment. (I) IMR90 ER:STOP and ER:RAS cells were transfected with control (NTP), *CASP1* or *CASP4*-targeting siRNAs. After 5 days of 4OHT treatment, caspase-4 proIL-1β and pRb were analyzed by immunoblotting. (J) Catalytically inactive (C258A) *CASP4* was overexpressed in IMR90 cells prior to LPS transfection (1 μg LPS/5 × 10^5 cells). 48 h after transfection, caspase-4 and pRb were analyzed by immunoblotting. (K) IMR90 cells were infected with an empty pRS vector (vector) or a pRS vector targeting either *CASP1* or *CASP4* prior to transfection with 0.1 μg LPS / 5 × 10^^5^ cells. *CCNA1*, *CDC6* and *BUB1* mRNA relative expression were quantified by RT-qPCR 48 h after LPS transfection. (L) IMR90 ER:STOP and ER:RAS cells were transfected with control, *CASP1*, *CASP4* or *GSDMD*-targeting siRNAs. BrdU incorporation was measured by IF 5 days after 4OHT addition. Statistical significance in A, G and L was calculated using two-tailed Student’s *t*-test. Statistical significance in C-F and K was calculated using one-way analysis of variance (ANOVA). ****P < 0.0001, \*\*\**P* < 0.001, \*\**P* < 0.01, and \**P* < 0.05. ns, not significant. All error bars represent mean ± s.e.m of 3 independent experiments.

---

**Table S1:**
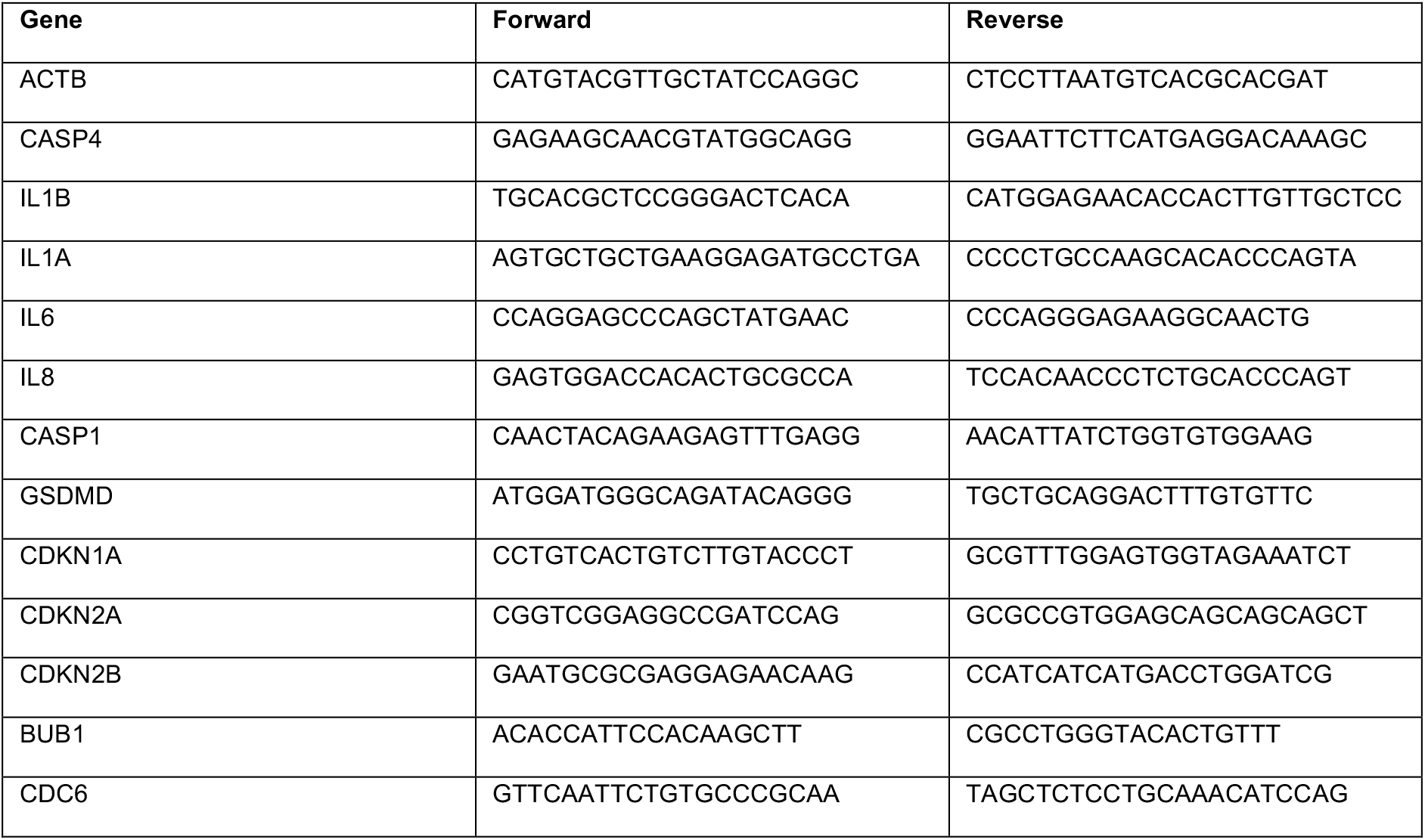
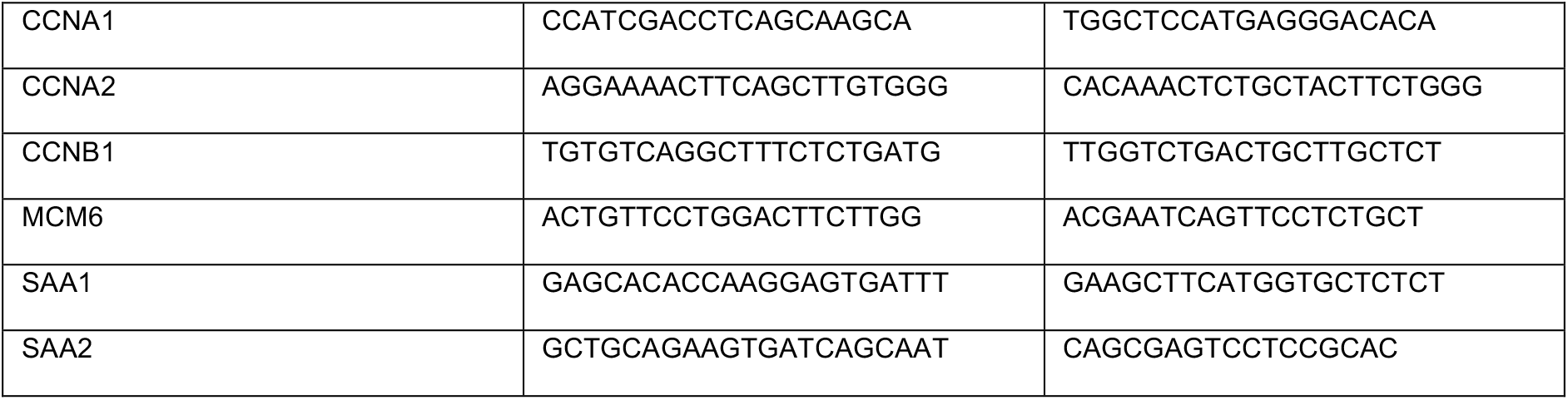
Primers used for mRNA gene expression analysis.

